# A small protein, derived from an alternative 53BP1 promoter transcript and expressed via translational reinitiation on an internal overlapping ORF, modulates proteasome activity

**DOI:** 10.1101/2021.08.23.457304

**Authors:** Inchingolo Marta Angela, Adamczewski Maxime, Humphreys Tom, Jaquier-Gubler Pascale, Curran Joseph Alphonsus

## Abstract

The complexity of the metazoan proteome is significantly increased by the expression of small proteins (<100 aas) derived from smORFs within lncRNAs, uORFs, 3’ UTRs and, more rarely, reading frames overlapping the CDS. These smORF encoded proteins (SEPs) can have diverse roles, ranging from the regulation of cellular physiological to essential developmental functions. We report the characterisation of a new member of this protein family, SEP^53BP1^, derived from a small internal ORF that overlaps the CDS that encodes 53BP1. Its expression is coupled to the utilisation of an alternative, cell-type specific, promoter coupled to translational reinitiation events mediated by a uORF in the alternative 5’ TL of the mRNA. The uORF-mediated initiation at the internal AUG^53BP1^ is conserved in metazoan species ranging from human to zebrafish. As such, it couples SEP^53BP1^ expression to the integrated stress response (ISR). We demonstrate that one function of this protein is to interact with, and stimulate, the activity of the 26S proteasome. As such, it opens the door to new approaches in the treatment of clinical conditions that arise due to the accumulation of toxic intracellular protein aggregates

## INTRODUCTION

Protein synthesis represents a key step in the regulation of gene expression. The differential recruitment of mRNA populations onto polysomes permits a rapid response to changes in the cellular environment. As such, it is a key process in the maintenance of homeostasis, and perturbations in its control are associated with numerous disorders. Translation can be subdivided into four main steps: initiation, elongation, termination and sub-unit recycling. Most regulation is exerted at initiation, and this has been confirmed in translational profiling studies covering the entire mammalian transcriptome ^1^. The ternary complex (TC) composed of Met-tRNAi-eIF2-GTP is first loaded onto the 40S ribosomal subunit in combination with a series of eukaryotic initiation factors (eIFs) to form the 43S pre-initiation complex (PIC). The PIC generally loaded onto the mRNA via the 5’ cap. Once recruited, it moves forward (5′→3′) scanning the mRNA 5’TL or UTR (Transcript Leader or UnTranslated Region) to locate the first AUG. The nucleotides flanking the AUG codon influence the efficiency of recognition, with the sequence 5’-ACCAUGG-3’ (the Kozak context: the nts in red being particularly important) being optimal^2^. If sub-optimal, scanning ribosomes will sometimes ignore the AUG codon and continue to the next. This phenomenon, known as leaky scanning, can produce N-terminal truncated proteins or proteins from overlapping reading frames^3, 4^.

The 5’TL contains a number of features that can regulate the translational readout during both PIC recruitment and subsequent scanning ^5^. This includes uAUGs and uORFs (upstream Open Reading Frames). Genomic analysis has estimated that ∼50% of human 5’TLs contain one or more uORFs^6, 7^. Both uAUGs and uORFs can function as translational repressors limiting PIC access to downstream start codons^8^. The amplitude of this repression is dictated by the uAUG context^9^. However, small uORFs (< 50 codons) can also couple the readout to stress and TC levels in the cell, via a process referred to as delayed reinitiation in which the 40S ribosome remains on the mRNA and continues to scan subsequent to translation of the uORF. This process permits access to start codons downstream of the AUG of the principle ORF (AUG^GENE^)^7, 10^. However, the efficiency of reinitiation at downstream start sites varies depending on parameters such as uORF length and the distance between the stop codon and the AUG. This process is conserved from human to zebrafish^11^. Reacquisition of the Met-tRNA by the 40S ribosome post-uORF termination is dependent on eIF2-GTP levels. When low, the slow reacquisition can cause a bypass of a proximal downstream AUG as the 40S is unable to re-recruit the TC necessary to form an initiation competent PIC. The TC levels respond to stress via the regulation of a series of “stress activated protein kinases” that include GCN2, HRI, PKR and PERK. These form the axe of the “integrated stress response” (ISR). Their substrate is the α subunit within eIF2.GDP generated during each round of translational initiation^12, 13^. Phosphorylation impedes GDP/GTP exchange and the subsequent TC regeneration. Thus, reinitiation in combination with leaky scanning offer the possibility to significantly increase the complexity of the mammalian proteome by permitting access to internal AUGs (iAUG).

Alterations in the 5’TL arise due to the use of alternative promoters (AP), transcriptional start site (TSS) heterogeneity and alternative splicing^14–16^, with studies suggesting that AP exceeds alternative splicing in generating transcriptome diversity^14^. A genome wide analysis revealed that ∼18% of human genes use multiple promoters^17^. Promoter switches change the nature of the first exon, and hence the 5’TL, and this event has been linked to a number of human pathologies^18, 19^. This switch is rarely complete but it can be amplified by the selective recruitment of one of the TL variants onto polysomes, as occurs with the MDM2 gene in tumour cells^20, 21^. Therefore, by generating 5’TL heterogeneity, which can be both tissue and cell type specific, alternative promoters regulate the protein readout, the proteome and ultimately the cellular phenotype^16^. Indeed, in a transcriptome/translatome analysis using a glioblastoma model, the authors concluded that selective polysomal recruitment of specific mRNA populations could initiate and drive tumour formation^22^.

We have discussed uORFs as translational regulatory elements. However, transcriptome analysis has identified thousands of yet non-annotated small open reading frames (smORFs) with the potential to encode biologically active peptides or SEPs (smORF-encoded proteins/peptides) smaller than 100 aas^23–26^. Detecting the products of smORFs, which are numerous and small, is technically not straightforward^27^. However, it has been facilitated by ribosome profiling^6^. This technique couples ribosome footprinting to high-throughput RNAseq and provides quantitative information about ribosome density across a transcript. It has been used to identify alternative START/STOP sites, initiation from non-AUG codons, translational pausing/frame-shifting as well as expression from uORFs and alternative ORFs^28^. A bioinformatics analysis of the ribosome profiling database revealed that 40% of long non-coding RNAs (lncRNAs) carry smORFs and are expressed in human cells^29^. Another source of SEPs are the uORFs, with ∼35% of mRNA coding genes having uORFs that are expressed^29^. These SEPs can act either *in-cis* to modulate downstream initiation events, or have distinct biological function(s)^7, 30^. Stalling of ribosomes over the AUG^GENE^ start site can also cause queuing of scanning ribosomes within the 5’TL. This can permit the expression of smORFs initiating on near-cognate codons with the potential to increase the SEP repertoire^31, 32^. Another interesting group are the cis-acting peptides that are responsive to environmental signals and have been coined “peptoswitches”^33^.

Both leaky scanning and reinitiation permit access to internal AUGs. When in-frame with the principle ORF this gives rise to N-terminally truncated proteins^34, 35^. When positioned internal and out-of-frame (ioORF: internal overlapping ORF), they represent a second source of smORFs. About 4% of human mRNAs appear to express SEPs from AUG codons downstream of the AUG^GENE^ ^29^. In fact, the expression of biologically active proteins from ioORFs has been known for some time. It was described in mammalian viral systems as far back as the early 1980’s^4, 36^. Nevertheless, its implications for the human proteome are only now beginning to be appreciated^37^. For example, within the ataxin-1 (ATXN1) transcript, a small ioORF starting 30nts downstream of the AUG^ATXN1^, and in the −1 reading frame, is expressed by leaky scanning^38^. The SEP, Alt-ATXN1, co-localises and interacts with Ataxin-1 within nuclear inclusions. The prion protein gene PRNP also expresses a novel polypeptide from an ioORF, referred to as AltPrP^39^. It localises at mitochondria, is up-regulated by ER stress and proteasomal inhibition and was detected in human brain homogenates, primary neurons, and peripheral blood mononuclear cells. Despite sizes smaller than 100 aas, the products of smORFs can have essential biological functions^23, 24^. In mice, the Mln smORF expresses a 46 aa SEP implicated in muscle contraction^40^. In humans, a 24 aa long SEP called humanin, synthesized from a lncRNA, is involved in apoptosis, interacting with BAX (Bcl-2-associated X protein)^41^, and the MRI-2 smORF (69 aas) has been implicated in DNA repair^24, 42^. Intriguingly, it has been proposed that in general, the expression of SEPs may be coupled to the stress response, an observation that would tie it in nicely with the process of translational reinitiation^43^. With regards to clinical medicine, a number of human cancer specific antigens are also derived from iORFs^44, 45^. Their expression reflects the change in the translational landscape that occurs with cellular transformation and they represent novel targets for immune based therapies^46^.

In this manuscript, we have extended on our earlier study in which we reported a differential RNAseq analysis on the tumoural MCF7 and non-tumoural MCF10 cell lines^47^. A number of genes were identified that exploited alternative promoters to generate 5’TL heterogeneity that could, in-turn, modulate the protein readout. One of these, the 53BP1 gene, uses two promoters (Fig. 1A). The P1 promoter (TSS12390) was active in both cell backgrounds. It generates two transcripts, referred to as V1 and V2 (NM_001141979.1, NM_001141980.1), which possess the same 5’TL but differ due to an alternative splicing event within the CDS (hereafter referred to as V1/2). The second P2 promoter (TSS20205) was more active in MCF7 cells^47^. It generates a V3 transcript (NM_005657.2) with a ∼278 nts 5’TL carrying a 5-codon uORF whose stop codon is 15 nucleotides upstream of the AUG^53BP1^ (Fig. 1A). We postulated, and now confirm, that this uORF directs reinitiation events at an ioORF that expresses a 50 aas SEP which we refer to as SEP^53BP1^. We provide evidence that uORF mediated expression of such a protein is conserved right through to zebrafish. The endogenous SEP^53BP1^ protein has been detected in a number of human cell lines of lymphoid origin and shows punctate staining in both cytoplasmic and nuclear compartments.

**Figure 1.**
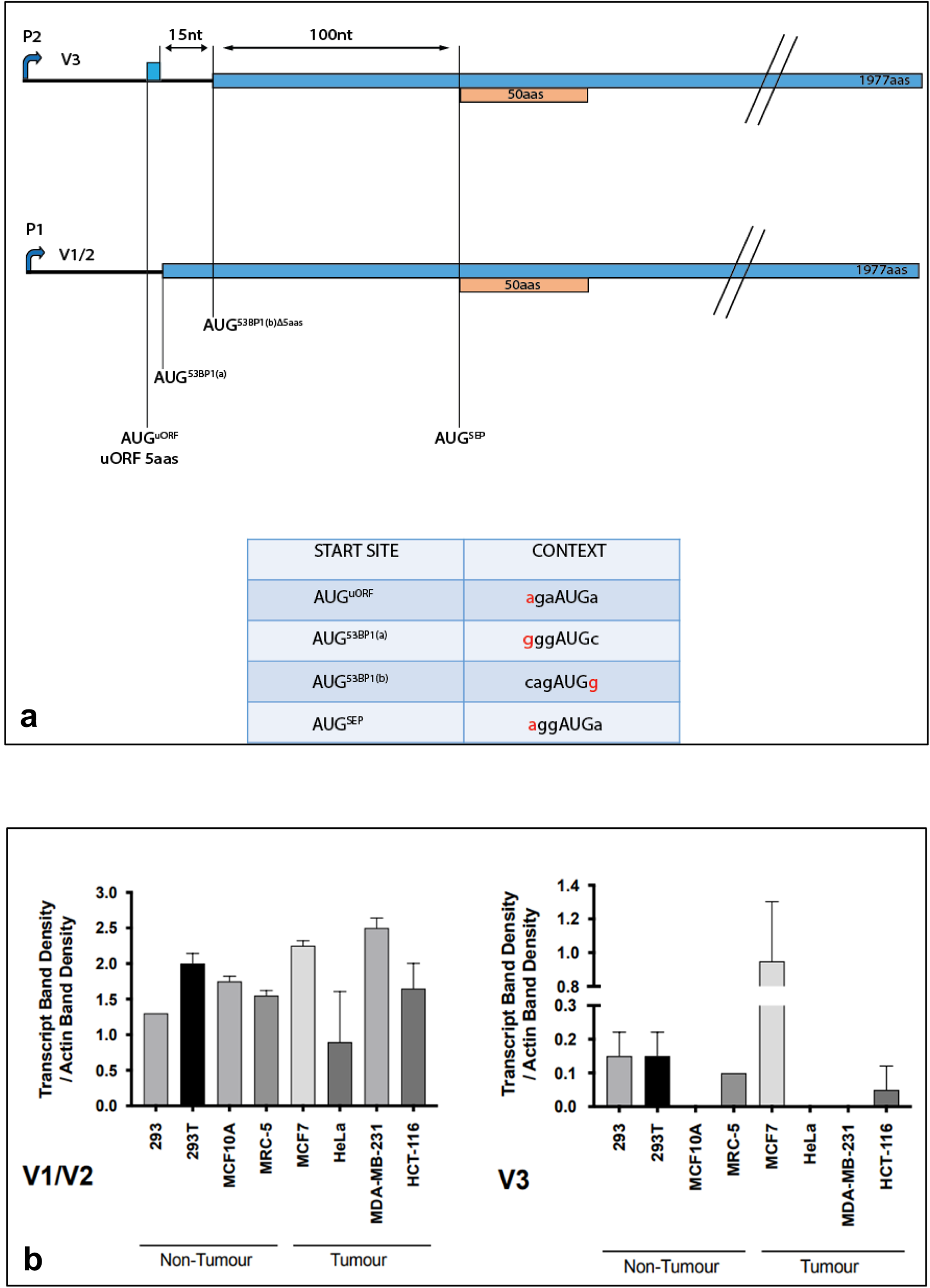
The human 53BP1 variants and their expression levels in tumoural and non-tumoural cell lines. (**a**) Schematic representation of the V1/2 and V3 transcript variants. P1 and P2 refer to the alternative promoters. The different AUG initiation sites are marked and there Kozak contexts are indicated in the lower table (nucleotides indicated in red indicate a positive context). The 53BP1 ORF is indicated in blue and the overlapping SEP ORF in orange. The small blue rectangle in V3 refers to the uORF. (**b**) Levels of each transcript variant in a range of cell lines. RT-PCR values for each variant were normalised to actin. The results are from biological triplicates. Bars indicate the SEM.

Interactome and biochemical studies indicate that it interacts with components of the proteasome machinery. Thus, we have identified a novel small protein whose expression is linked to a promoter switch, coupled to a stress responsive translational reinitiation event on an AUG^ioORF^. At least one function of the expressed SEP protein is to modulate proteasome activity. We discuss these observations in the light of current models for SEP function, and the potential therapeutic applications of increased SEP^53BP1^ expression in the treatment of human neurodegenerative diseases^48^.

## RESULTS

### Organisation and expression profiles of the 53BP1 gene transcripts

In our earlier work, we reported on a differential RNAseq analysis comparing the tumoural MCF7 and non-tumoural MCF10 cell lines^47^. The 53BP1 gene was a particularly intriguing hit. It uses two promoters. The P1 promoter is active in both cell backgrounds. It generates two transcripts, named V1/2, originating from alternative splicing but carrying the same 5’TL. The mRNA has two potential AUG start codons in the 53BP1 ORF, located at the end of the first and beginning of the second exons, and separated by four codons, hinting at two N-terminal isoforms (Fig. 1a). These we refer to as AUG^53BP1(a)^, which has a relatively good Kozak context, and AUG^53BP1(b)^ whose context is poor (Fig. 1a, lower panel). The 5’TL is ∼113nts long, 71% G/C and contains no uAUGs. The second P2 promoter was more active in MCF7 cells (Fig. 1b)^47^. Based upon CAGE analysis it generates a V3 transcript with a ∼278 nts 5’TL carrying a 5 codon uORF whose stop codon is 15 nucleotides upstream of the AUG^53BP1(b)^ (Fig. 1a: AUG^53BP1(a)^ is in the first exon of the V1/2 transcript). Luciferase based reporter assays revealed that the V3 5’TL was more repressive than V1/2 with regards to initiation events on the AUG^53BP1^ start sites, due to the uORF^47^. Furthermore, polysome gradient profiling of the two cell lines revealed that whereas the V1/2 transcript was mainly polysomal in both, the V3 transcript was polysomal only in the tumoural MCF7 cells^47^. Therefore, at the outset of our current study we evaluated to what extent P2 promoter activity was a marker of the tumoural phenotype. We performed an RT-PCR analysis of V1/2 and V3 across a range of established tumoural and non-tumoural cell lines available in the lab. No clear correlation with the tumoural phenotype was observed (Fig. 1b).

### The protein readout from the V3 mRNA is different

The small uORF in the V3 5’TL could promote delayed reinitiation events downstream of the AUG^53BP1^. Examination of the human sequence reveals that the next start codon downstream AUG^53BP1(b)^ opens an ioORF, +1 relative to the 53BP1 ORF, that would encode a polypeptide of 50 aas that we named SEP^53BP1^ (Fig. 1a). To monitor expression in the V1/2 and V3 5’TL backgrounds at both the AUG^53BP1^ and AUG^SEP^, we inserted the sequences upstream of the AUG^SEP^ into our LP/SP overlapping ORF reporter^10, 49^. This fuses the 53BP1 ORF to LP (which carries a FLAG and HA tag) and the AUG^SEP^ to SP (which carries a MYC and HA tag: the AUG^SEP^ and its Kozak context were retained) (Fig. 2a). It allows us to follow initiation events at the AUG^53BP1^ (we were unable to distinguish between the sites AUG^53BP1(a)^ and AUG^53BP1(b)^ on V1/2: however, based upon context we presume that the former is the major start codon) (Fig. 2a). Transient expression assays in HEK293T cells, revealed that the V1/2 5’TL directed initiation events mainly at AUG^53BP1^ whereas with V3 the majority of initiation events occurred at AUG^SEP^ (Fig. 2a). This pattern was also observed in transient assays performed in MCF10 and MCF7 cells (Fig. 2b). To monitor the impact of the V3 uORF on the readout we mutated its stop codon (UGA→UGC: V3^UGA/UGC^) thereby fusing the uAUG to the 53BP1 ORF (Fig. 2b). This effectively removes events arising from delayed reinitiation. When transiently expressed in HEK293T, MCF10 and MCF7 cells, the V3^UGA/UGC^ directed expression mainly from the AUG^uORF^ (Fig. 2b). This would be consistent with its good Kozak context (Fig. 1a). Leaky downstream scanning to the AUG^53BP1^ and AUG^SEP^ was weak in the HEK293T and MCF10 cell backgrounds but more evident in MCF7 cells.

**Figure 2.**
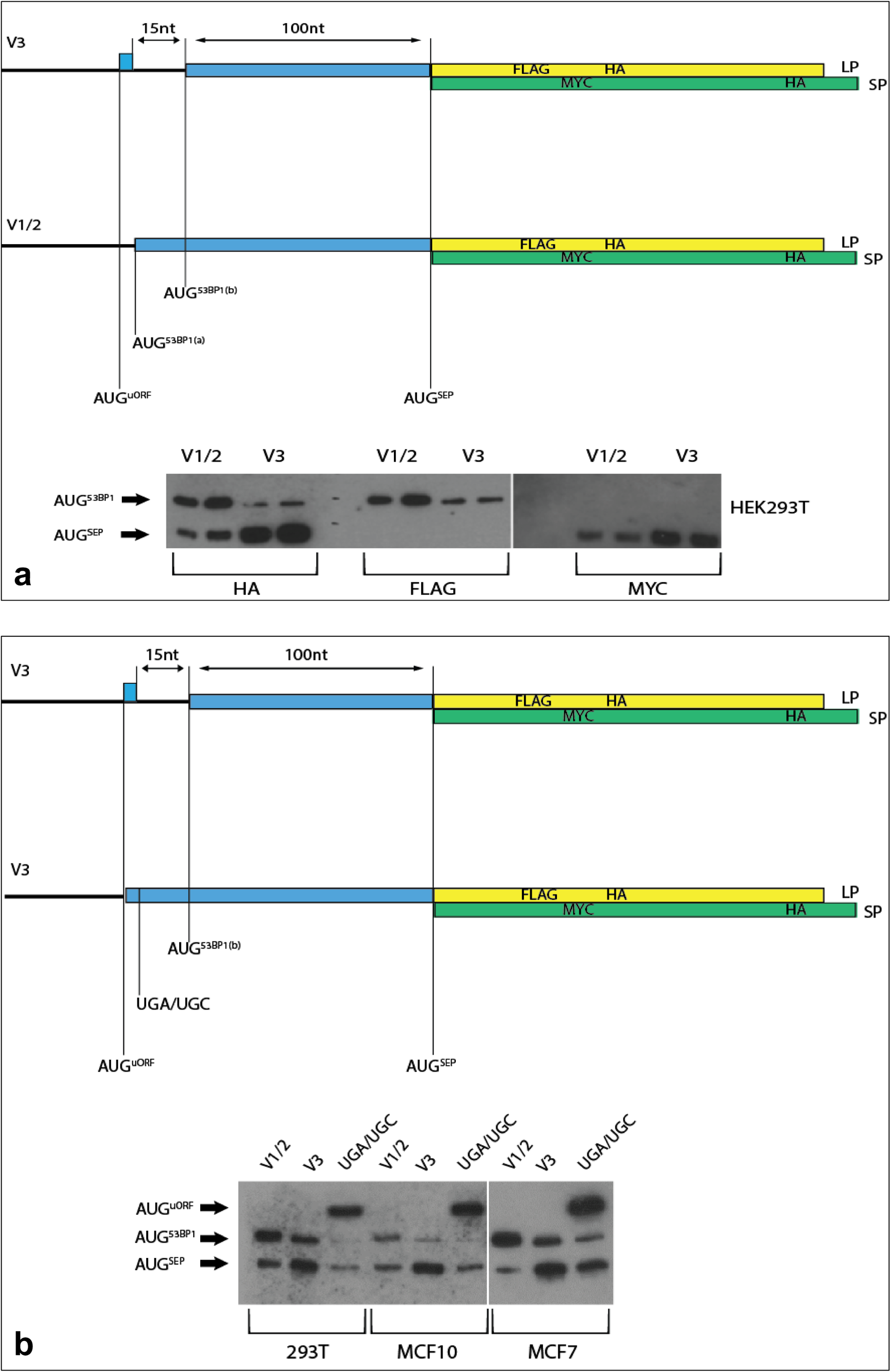
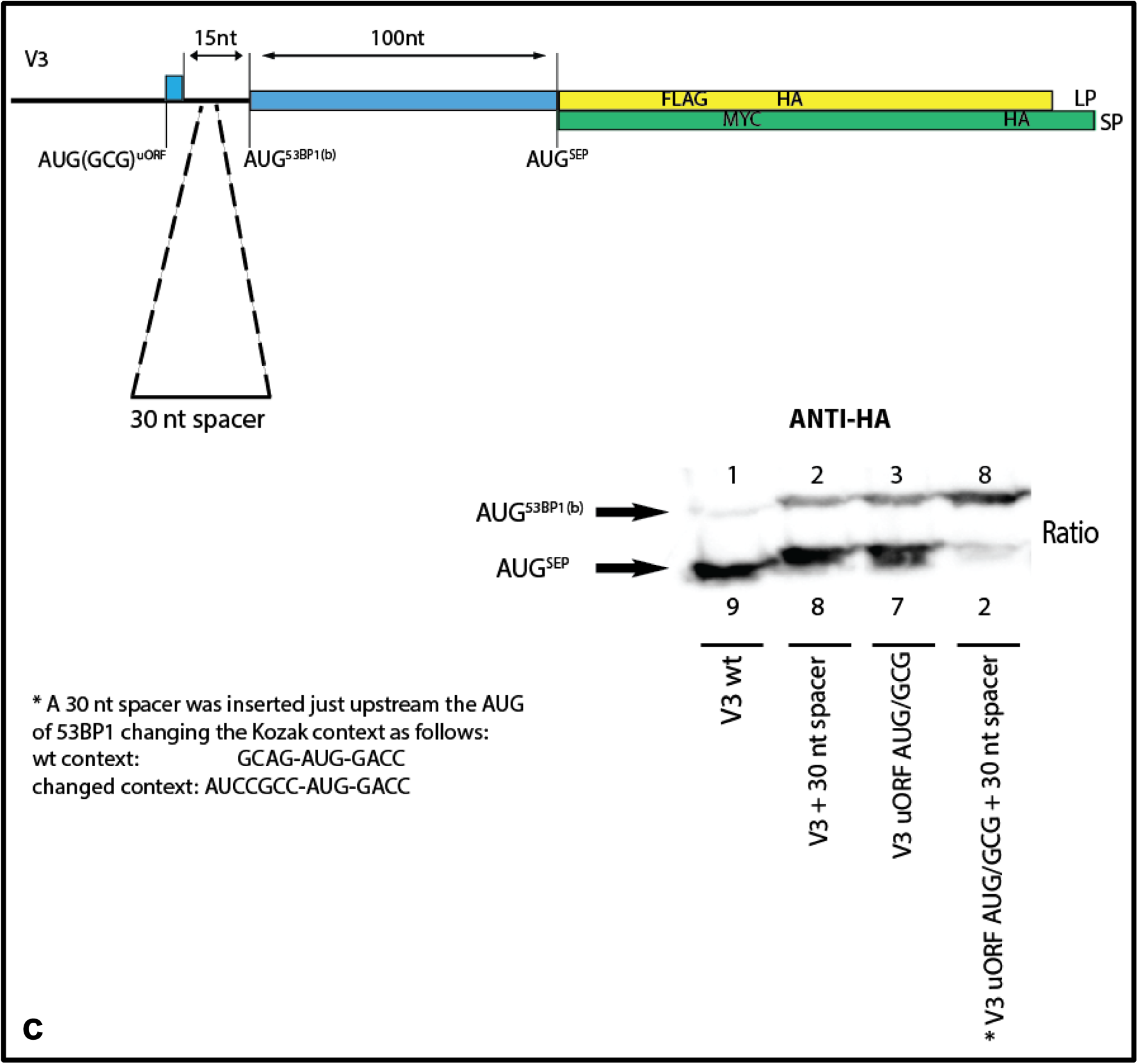
Reporter assays monitoring start site selection on the 53BP1 variants. (**a**) Upper panel: Schematic representation of the LP/SP reporter fused to the V3 and V1/2 sequences upstream of the AUG^SEP^. The LP ORF (yellow rectangle) carries FLAG/HA tags and was fused in-frame to the 53BP1 ORF (indicated in blue). The AUG^53BP1^ started the SP ORF (green rectangle) which carries MYC/HA tags. These constructs were transiently expressed in HEK293T cells and protein steady-state levels were monitored on Western blots using the anti-HA, anti-FLAG and anti-MYC Abs (lower panel). (**b**) Upper panel: Schematic representation of the V3 construct with the UGA/UGC mutation that fused the uORF to the 53BP1 ORF. The V1/2, V3 and V3UGA/UGC constructs were transiently expressed in HEK293T, MCF10A and MCF7 cells and steady-state protein levels were monitored on Western blots using the anti-HA Ab which monitors expression from both overlapping ORFs (lower panel). (**c**) A schematic representation of the 30nt spacer introduced between the uORF stop codon and the AUG^53BP1^. This construct, and a series of other constructs, as listed in the lower panel, were transiently expressed in HEK293T cells and expression monitored by Western using the anti-HA Ab. The numbers indicate the intensity of the bands normalised to the AUG^53BP1(b)^ signal that was set as 1.

The results demonstrate high levels of initiation at the AUG^SEP^ in transcripts carrying the V3 5’TL and this is mediated by its small uORF. To confirm a role for delayed reinitiation we introduced a 30 nts spacer element between the uORF and the AUG^53BP1(b)^. Consistent with a reinitiation model, this increased initiation events at the AUG^53BP1(b)^ relative to AUG^SEP^ (Fig. 2c: compare lanes 1 and 2). The expression pattern observed with the 30 nts spacer construct was similar to that obtained when the uAUG was changed to GCG (Fig. 2c: compare lanes 2 and 3). In the uAUG/GCG mutant, we still observed significant initiation events at the AUG^SEP^. That this arose due to leakiness of the AUG^53BP1(b)^ was confirmed by introducing changes that improved its context (..CAGAUGG… → …GCCAUGG) (Fig. 2c: compare lanes 3 and 4). In conclusion, the presence of a short uORF positioned close to a very leaky downstream AUG^53BP1(b)^ means that the major initiation events on the V3 transcript take place on AUG^SEP^. This would direct the expression of a smORF-encoded peptide (SEP) of 50 aas (SEP^53BP1^)^23, 24^.

### The configuration of the V3 5’TL and the ioORF are conserved

Sequence alignment suggests that the SEP^53BP1^ ioORF is conserved across vertebrates and can be found even in zebrafish (Danio rerio) (Fig. 3a). Likewise, uORFs are frequently found within the annotated 5’TLs of the 53BP1 gene. For example, zebrafish have a single promoter expressing a single 5’TL variant with an uORF of 19 codons (GenBank: BC129236.1 and NM_001080170: longer than the human) whose stop codon is 15 nts upstream of the first AUG^53BP1a^ (similar to human) (Fig. 3b). uORFs are frequently present in zebrafish transcripts and, as in mammals, they serve to modulate the translational readout^11^. The zebrafish ioORF would express a SEP polypeptide of 40 aas (Fig. 3a). The nucleotide spacing between the AUG^53BP1a^ and AUG^ioORF^ is 400 nts in zebrafish compared to 97 nts in the human V3 mRNA. Within this 400 nt region there is a second AUG in the 53BP1 ORF (AUG^53BP1b^) that could express an N terminally truncated (Δ121 aas) 53BP1 protein (Fig. 3B). To examine initiation events on this transcript, we RT-PCR cloned all 53BP1 sequences upstream of the ioORF STOP codon (changing it at the same time to a sense codon) starting from total zebrafish embryonic RNA. This was then fused to our LP/SP reporter to generate 53BP1ZLP/SP WT (Fig. 3B). To monitor the role of the uORF on start site selection, a number of mutations were created. The uORF^AUG/GCG^ removed the start codon and the uORF^UAA/AGG^ fused the uORF to the 53BP1/LP ORF in the reporter (Fig. 3b). We also exploited two BamHI sites, one positioned just before the uORF UAA stop codon and the second just after the AUG^53BP1a^. Deletion of the small BamHI fragment removed both the uORF^UAA^ and AUG^53BP1a^ codons fusing uORF to the ORF of 53BP1/LP (Fig. 3b: 53BP1ZLP/SPΔBam). As in the human reporter construct, the ioORF was fused to the SP reading frame (Fig. 2). In the WT background, we could detect products from AUG^53BP1a^, AUG^53BP1b^ and AUG^ioORF^, with the latter corresponding to the zebrafish SEP^53BP1^ (Fig. 3c, lane 2). Removal of the uAUG significantly enhanced expression at the AUG^53BP1a^ but did not impact significantly on the downstream start sites (Fig. 3c, lane 3). These latter initiation events would now arise due to leaky scanning through AUG^53BP1a^ whose context is poor (Fig. 3b). Thus as in humans, the uORF in zebra fish represses 53BP1 expression. Fusing the uORF to the 53BP1 ORF, either by the uORF^UAA/AGG^ mutation (Fig. 3c, lane 4) or the ΔBamH1 deletion (which also removes AUG^53BP1a^: Fig. 3c, lane 5) produced a single band on the blot whose slower migration indicates that it arises from an initiation event on AUG^uORF^. The “non-leakiness” of this start codon would be consistent with its good Kozak context (Fig. 3b). We confirmed this by introducing the uORF^AUG/GCG^ mutation into the ΔBamH1 background. The slow migrating band was lost and we restored the expression of products from the AUG^53BP1b^ and AUG^ioORF^ (Fig. 3c, lane 7). Thus, in zebrafish the uORF is also permitting initiation events downstream of the AUG^GENE^ (in this case AUG^53BP1a^). These downstream initiation events can give rise to N-terminal truncated forms of the 53BP1 protein and the expression a SEP^53BP1^. However, unlike the human V3 transcript the single zebrafish 5’TL assures robust expression from all initiation sites.

**Figure 3.**
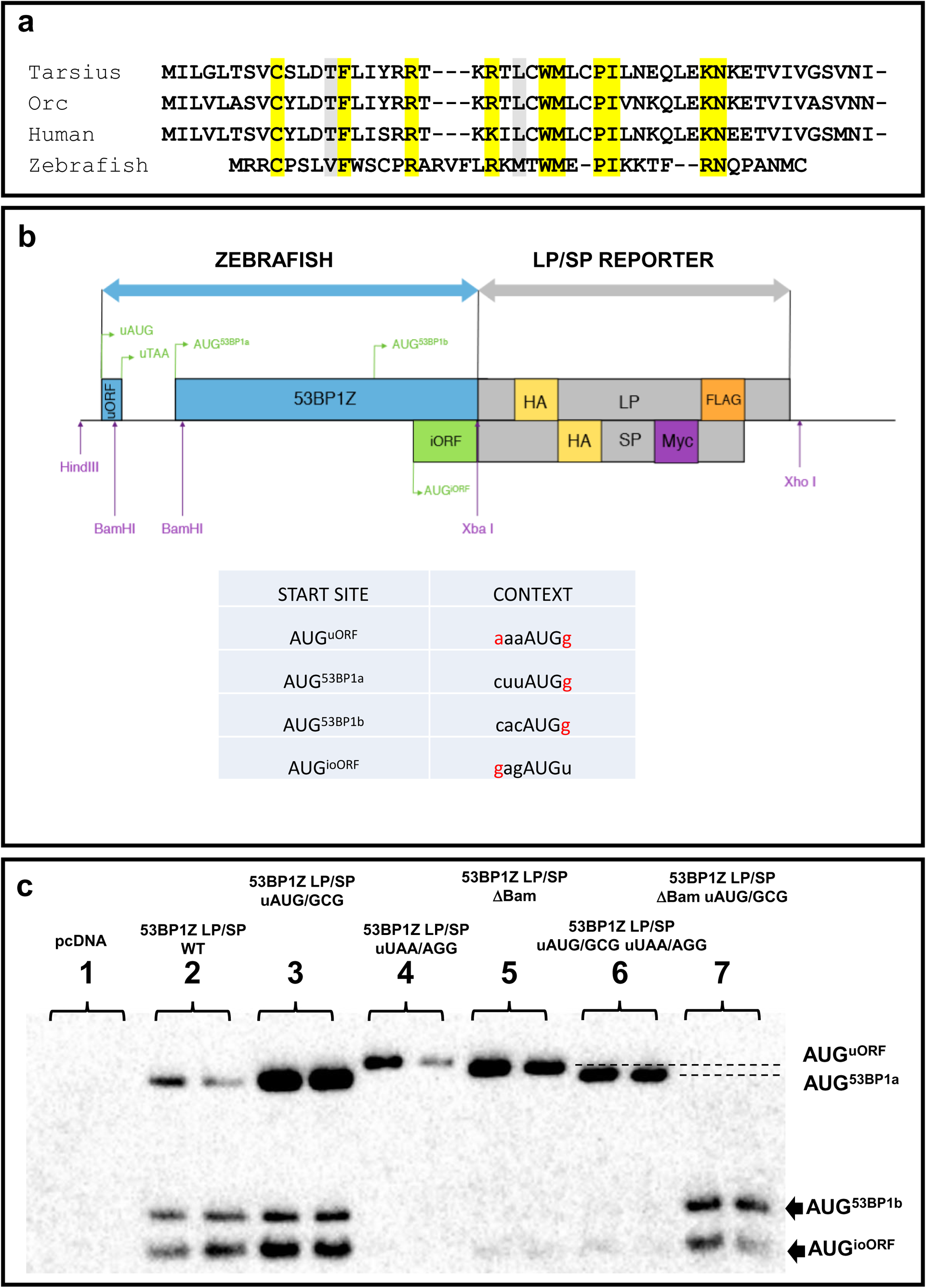
Delayed reinitiation mediated by a uORF promotes expression at the AUG^SEP^ also in Danio rerio (zebrafish). (**a**) Alignment of SEP^53BP1^ proteins in a range of vertebrates. Highly conserved amino acids across all four species are highlighted in yellow and those moderately conserved in grey. (**b**) Schematic representation of the LP/SP reporter construct generated to monitor initiation events on the zebrafish transcript. Sequences downstream of the iORF stop codon (which was removed) were fused to the reporter, with the 53BP1 ORF (blue rectangle) fused to LP and the ioORF (green rectangle) fused to SP. The position of the HA, FLAG and MYC tags are all indicated. The lower panel indicates the Kozak context for all the AUG initiation codons. Favourable nucleotides are highlighted in red. (**c**). A series of LP/SP zebrafish constructs (as indicated above each lane) were transiently expressed in HEK293T cells. Western blots were performed with the anti-HA Ab. Each transfection was performed in duplicate.

### Studies on the human SEP53BP1

Polyclonal Abs against the SEP^53BP1^ were generated using two peptides that spanned most of the protein (VLTSVCYLDTFLISRRTKKILC and WMLCPILNKQLEKNEETVIVGC: Proteogenix, France). The Ab did not detect SEP^53BP1^ expression in HEK293T cells (Fig. 4a, lane 2), an observation that would be consistent with the low levels of the V3 transcript in this cell line (Fig. 1b). We therefore generated an authentic full-length V3-53BP1 cDNA clone and transiently expressed it in HEK293T cells (Fig. 4a). A doublet band co-migrating with the SEP^53BP1^ protein expressed in a WGE (the *in-vitro* system was programmed with a capped/poyladenylated mRNA covering the SEP^53BP1^ ORF) was detected on blots (Fig. 4a, lanes 1 and 3). Proof that this arose from initiation events at the SEP^53BP1^ AUG codon came from both altering the AUG^SEP53BP1^ Kozak context from good to bad (…agg**AUG**a…→…cgg**AUG**a…), which impacted negatively on expression (Fig. 4a, lane 4), and changing the AUG^SEP53BP1^ to GCG which ablated all expression (Fig. 4a, lane 5).

**Figure 4.**
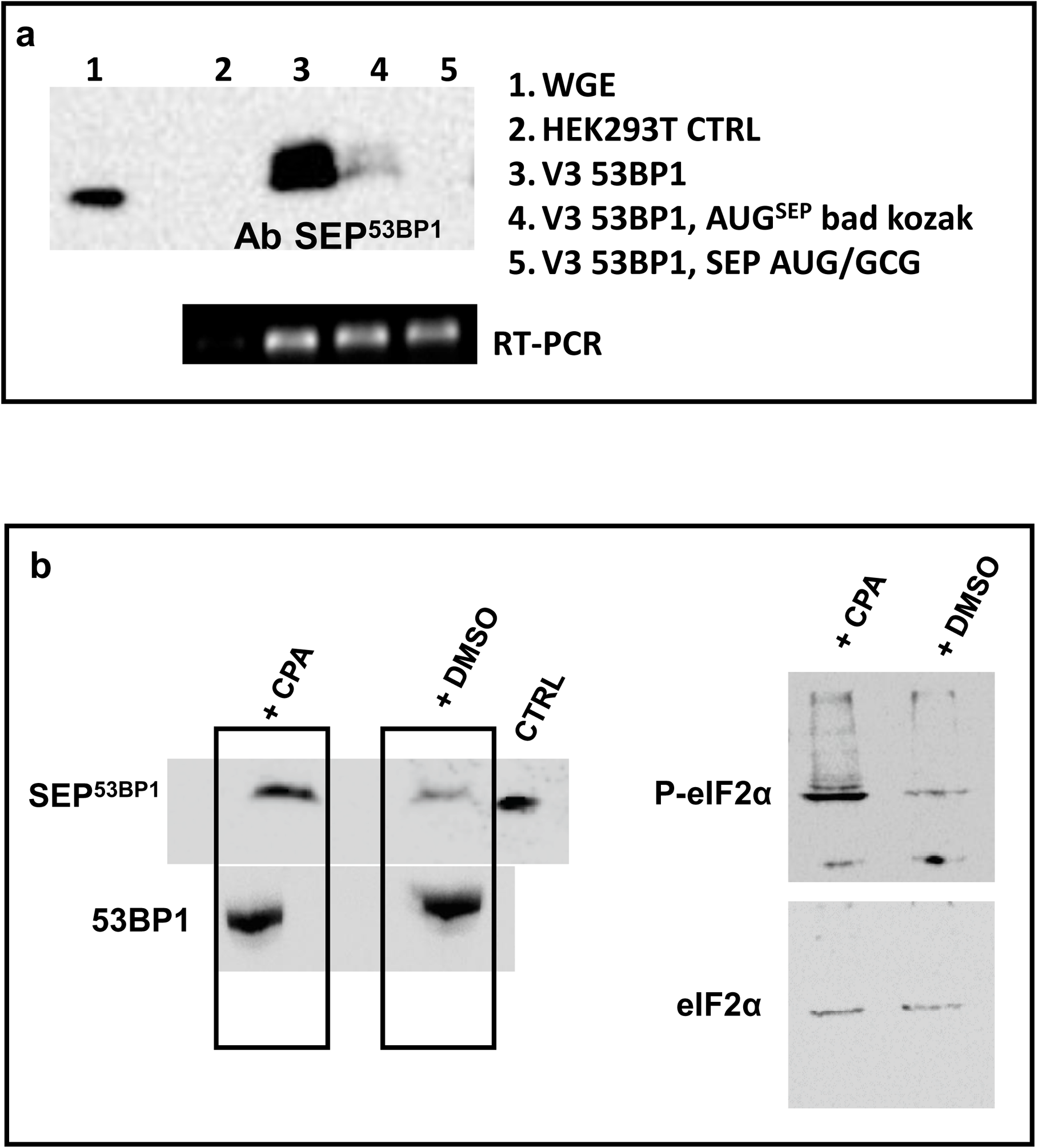
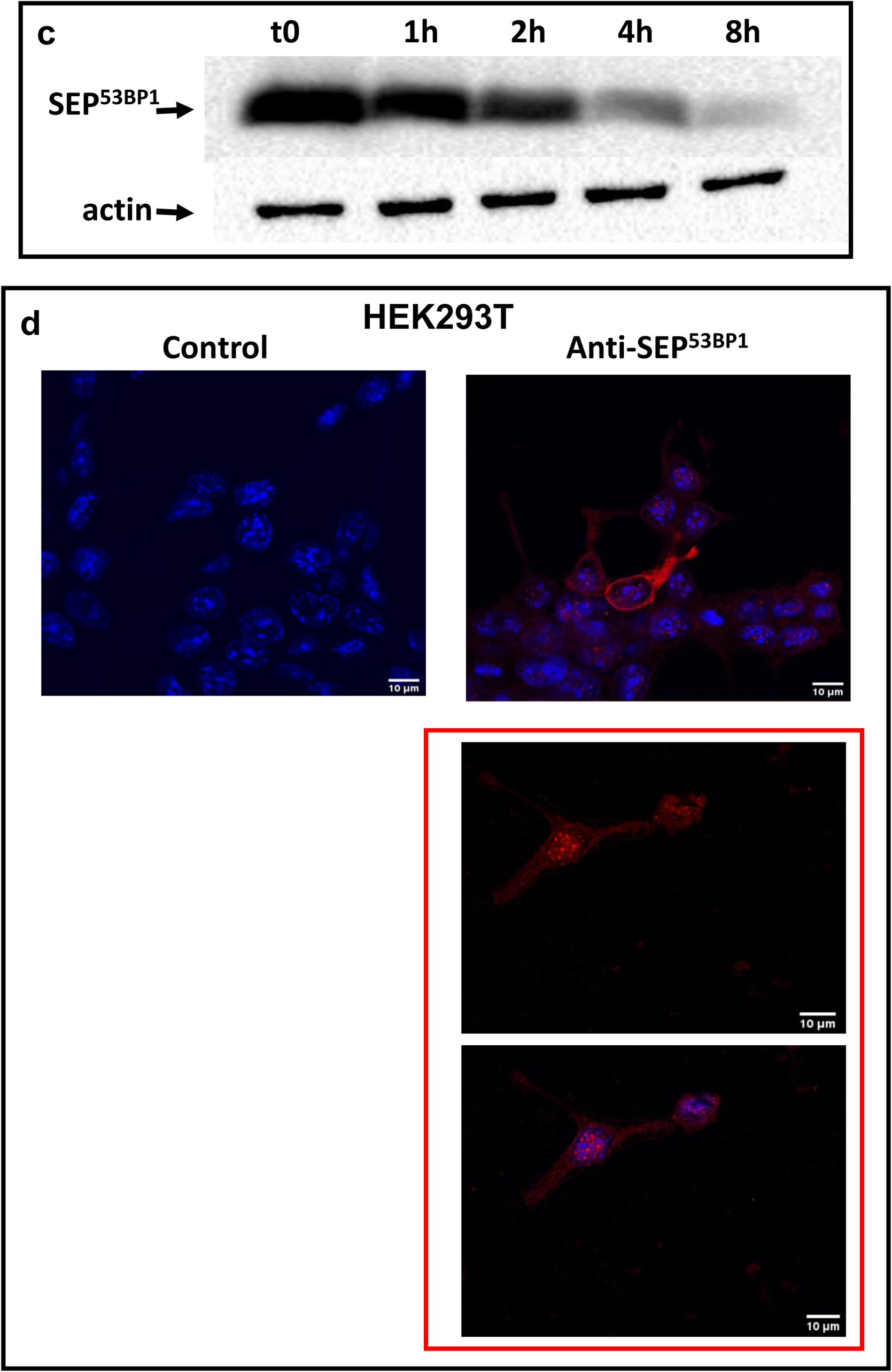
SEP^53BP1^ expression is increased during cellular stress. (**a**) A full-length human V3 53BP1 clone was generated and transiently expressed in HEK293T cells (lane 3). SEP^53BP1^ expression was monitored by Western blotting using an anti-SEP^53BP1^ polyclonal Ab. As a marker, an *in-vitro* generated SEP^53BP1^ mRNA was translated in a WGE (lane 1). Confirmation that the band observed in the transfected cells arose from initiation events at the AUG^SEP^ came by weakening the Kozak context (lane 4) and changing the AUG→GCG (lane 5). Transcript expression levels were monitored by RT-PCR (lower panel). (**b**) The V3 53BP1 clone was transiently expressed in HEK293T cells treated with either DMSO or the drug CPA for 16 hrs. SEP^53BP1^ and 53BP1 levels were detected by Western blotting (right hand panels: CTRL indicates a lane in which SEP^53BP1^ was expressed in WGE). Cellular stress was monitored by blotting using Abs against eIF2α and phospho-eIF2α (P-eIF2α: right-hand panels). (**c**): SEP^53BP1^ was transiently expressed in HEK293T cells seeded in a six well plate. At 16 hrs post-transfection cells were treated with cycloheximide (100 μg/mL) and recovered at the times indicated in the panel. The levels of SEP^53BP1^ and actin were monitored by immunoblotting. (**d**) IF studies using HEK293T cells transiently expressing SEP^53BP1^. The control was performed without the primary Ab. Nuclei were stained with DAPI (blue). The insert below the Anti-SEP^53BP1^ panel shows cells in which staining was mainly nuclear.

Furthermore, and consistent with the reinitiation model derived from the reporter assays (Fig. 2), expression increased under stress conditions that activated PERK (treatment with the drug Cyclopiazonic acid, CPA), thereby increasing the intracellular phospho-eIF2α levels (Fig. 4b)^50^. The transiently expressed SEP^53BP1^ had an intracellular half-life of 2.5 hrs (Fig. 4c). Immunofluorescence (IF) imaging of transfected HEK293T cells revealed a mainly cytoplasmic localisation (Fig. 4d). However, staining could be observed in the nucleus and, in rare occasions, it was almost exclusively nuclear (Fig. 4d, lower panel).

### Detection and localisation of the endogenous SEP53BP1 protein

Transient expression assays have allowed us to elucidate the mechanism by which P2 promoter activation will permit the expression of a novel SEP. However, at this point in the study it was necessary to detect the endogenous protein, and determine the cellular compartment(s) in which it accumulates as a route towards function. We had already observed that P2 promoter activity and V3 transcript levels are regulated in a cell-specific manner (Fig. 1b). Furthermore, we had reported that polysomal recruitment of V3 could also be cell specific^47^. With this in mind, we scanned the ribosome-profiling database (http://sysbio.sysu.edu.cn/rpfdb/index.html). The image in Supplementary Fig. 1 was extracted from a study performed by the Brosch lab using THP-1 cells (a human acute monocytic leukaemia cell line: https://www.ncbi.nlm.nih.gov/geo/query/acc.cgi?acc=GSE39561)^51^. The accumulation of reads around both the AUG^53BP1^ and AUG^SEP53BP1^ would be consistent with their utilisation as start sites. We therefore performed polysomal analysis of the total, V1/2 and V3 mRNAs in this cell background (Fig. 5a). Only a minor fraction of the total 53BP1 gene transcripts were polysomal (26%) (Fig. 5a, left hand profile). Concerning V3, very little was associated with light polysomes (9%) although a more significant fraction was observed within the heavy polysomes (44%), in particular the heaviest fraction (fraction 11, 32%). It is worth remembering that the ioORF is only 150 nts in length and can accommodate a maximum of five elongating ribosomes^6, 52, 53^. This would mean that V3 transcripts in the heavy polysomal fraction, that we define as >5 ribosomes per transcript, must be translating both ORFs. We also analysed another lymphocyte cell line that was available to us, namely Raji cells (Fig. 5a, right hand profile: no ribo-profiling data is available for this cell line). The polysomal profiles indicated that the majority (80%) of the 53BP1 transcripts were polysomal and this was also observed with both V1/2 (78%) and V3 (87%) (Fig. 5a). Immunoblots detected SEP^53BP1^ expression in both these cell lines but only weak expression in MCF7 cells, the cell line in which we originally reported V3 expression (Fig. 5B)^47^. Curiously, in THP-1 cells, slower migrating bands were also detected reminiscent of the doublet observed with the transiently expressed protein (Fig. 4b), suggesting that post-translational modifications may be occurring. Confocal imaging of the endogenous protein in both THP-1 and Raji cells revealed punctate staining in both nuclear and cytoplasmic compartments (Fig. 5c). Z-stacking analysis confirmed the presence of endogenous SEP^53BP1^ in the nucleus (see Supplementary animation 1 and 2 for THP-1 and Raji, respectively).

**Figure 5.**
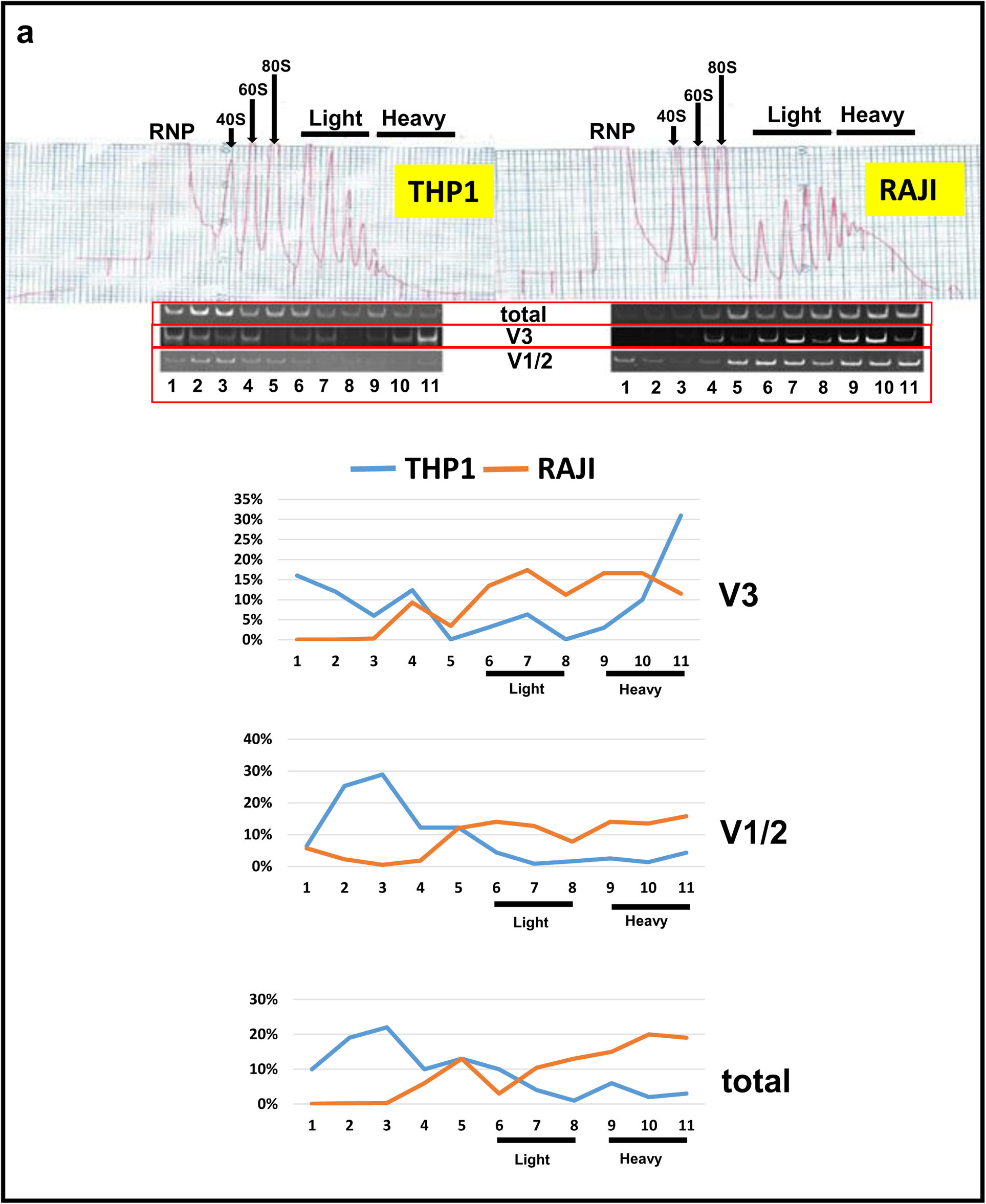
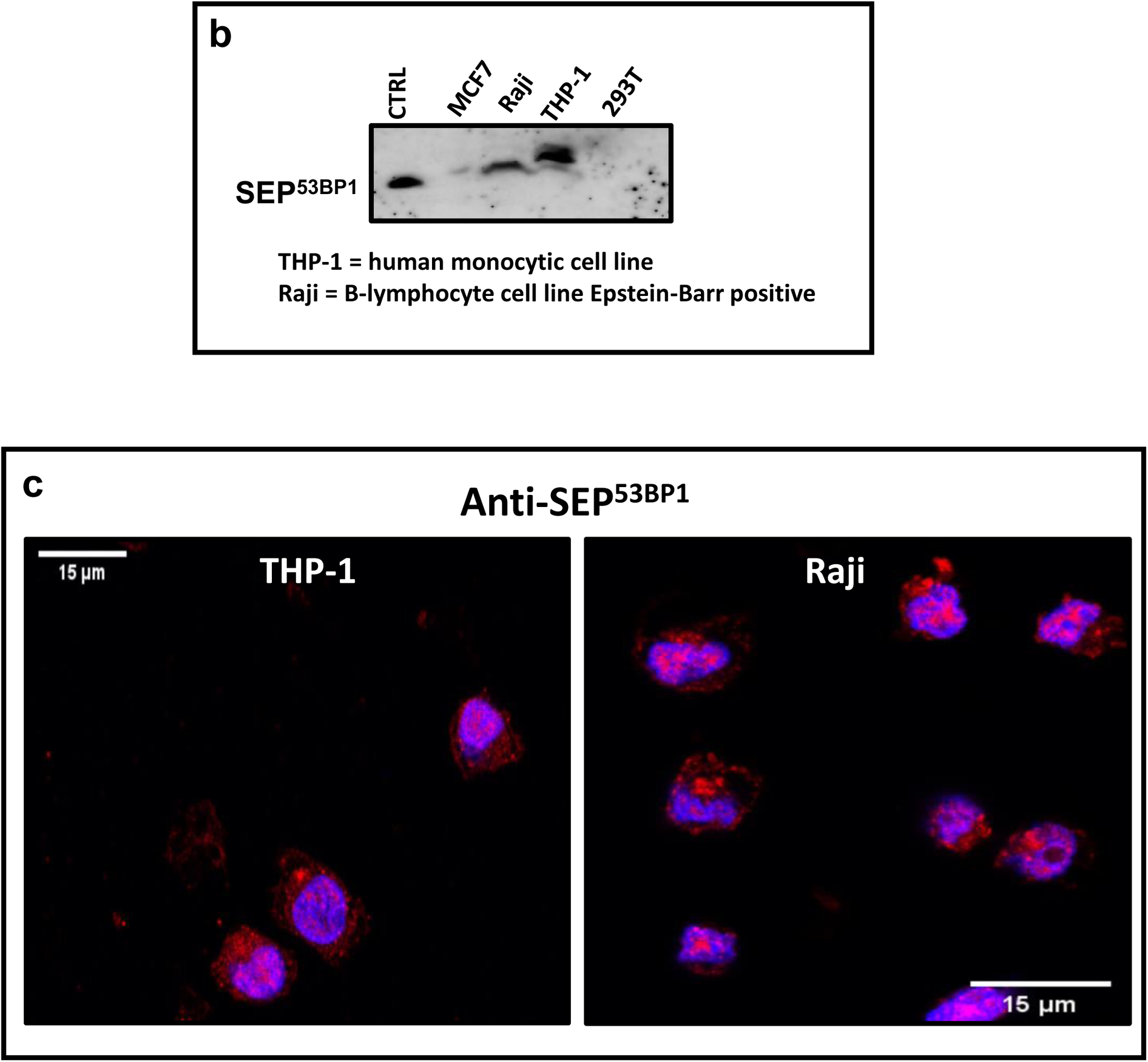
Detecting the endogenous SEP^53BP1^ protein. (**a**): Upper panel: Polysome gradient profiles from THP-1 and Raji cells indicating the RNP (ribonucleoprotein), 40S, 60S, 80S, light and heavy polysomal fractions. Gradients were collected as eleven equal fractions. RT-PCR was performed on equal amounts of RNA from each fraction using primer sets specific for each variant and the total 53BP1. Amplicons were resolved on polyacrylamide gels (panels below the profiles), quantitated and plotted graphically as a percentage of the total signal for each variant (lower images). (**b**) Approximately 3×107 cells (MCF7, THP-1, Raji and HEK293T) were lysed in CSH buffer and immunoprecipitated with anti-SEP^53BP1^ Ab. Immune-complexes were recovered on Protein-G magnetic beads and analysed by Western blotting using the same Ab. The CTRL is the SEP^53BP1^ protein expressed in WGE. (**c**) IF in THP-1 and Raji cells using the SEP^53BP1^ Ab (red) superimposed on the cell nuclei (blue). Images were generated by confocal microscopy.

### The SEP53BP1 interactome

To gain insights into function we employed a yeast-2-hybrid (Y2H) screen to identify partners. SEP^53BP1^ was used as a prey, and screened against a peptide library generated from a human B cell Lymphoma_RP1. This background was selected because we had observed SEP^53BP1^ in two cell lines of lymphoid origin. Around 51 million interactions were tested and 5 genes (PSMA7, UBQLN4, TRIP12, MAPRE1, BCOR) gave interactions with good confidence levels (Fig. 6a). The selected interaction domain (SID) for each prey is depicted in Supplementary Fig. 2. String analysis connected three of the five genes (PSMA7, UBQLN4, TRIP12) to proteasome biology (Fig. 6a). We sought to biochemically validate this analysis focusing on the protein products of the first two genes. PSMA7 encodes the α4 subunit of the 20S proteasome barrel and it plays a key role in its assembly^54, 55^. Sedimentation analysis performed on cytoplasmic extracts prepared from HEK293T cells transiently expressing SEP^53BP1^ revealed that a fraction of the 50 amino acid protein co-sedimented with the α4 protein in fractions 4-6 (Fig. 6b, upper panel). Extracts contained ATP to ensure 26S proteasome integrity during the assay^56, 57^.That these fractions corresponded to the 20S/26S proteasome was confirmed by disruption of the complexes using SDS treatment of the extracts prior to gradient loading. (Fig. 6b, lower panel). It should be noted that whereas only a fraction of SEP^53BP1^ co-sedimented with α4 in native conditions, the majority of the protein entered into the gradient, and a significant fraction was found in the pellet ribosomal fraction (as indicated by the presence of the ribosomal protein S6: Fig. 6b, upper panel). However, our interactome study did not give any hits with components of the translational machinery. Overall, the sedimentation profile of SEP^53BP1^ is quite remarkable considering its small molecular size (very little of the protein remains on the top in fraction 10). These results suggest that it has multiple interacting partners in the cell. We directly demonstrated the α4-SEP^53BP1^ association by co-IP performed on HEK293T cell extracts transiently expressing the latter (Fig. 6c) and this corroborated their intracellular co-localisation (Fig. 6d). UBQLN4 also plays a role in the regulation of intracellular protein degradation by mediating the proteasomal targeting of misfolded or accumulated proteins^58^. Its over-expression, as observed in some human tumours, also represses homologous DNA repair^59^. It did not enter into our glycerol gradients, suggesting that under our assay conditions the protein was mainly free in the extracts (Fig. 6b), and we were unable to co-IP transiently expressed SEP^53BP1^ by pulling down the endogenous UBQLN4 protein. Nonetheless, an interaction could be demonstrated by co-IP in cells transiently expressing both UBLQN4 and SEP^53BP1^ (Fig. 6e).

**Figure 6.**
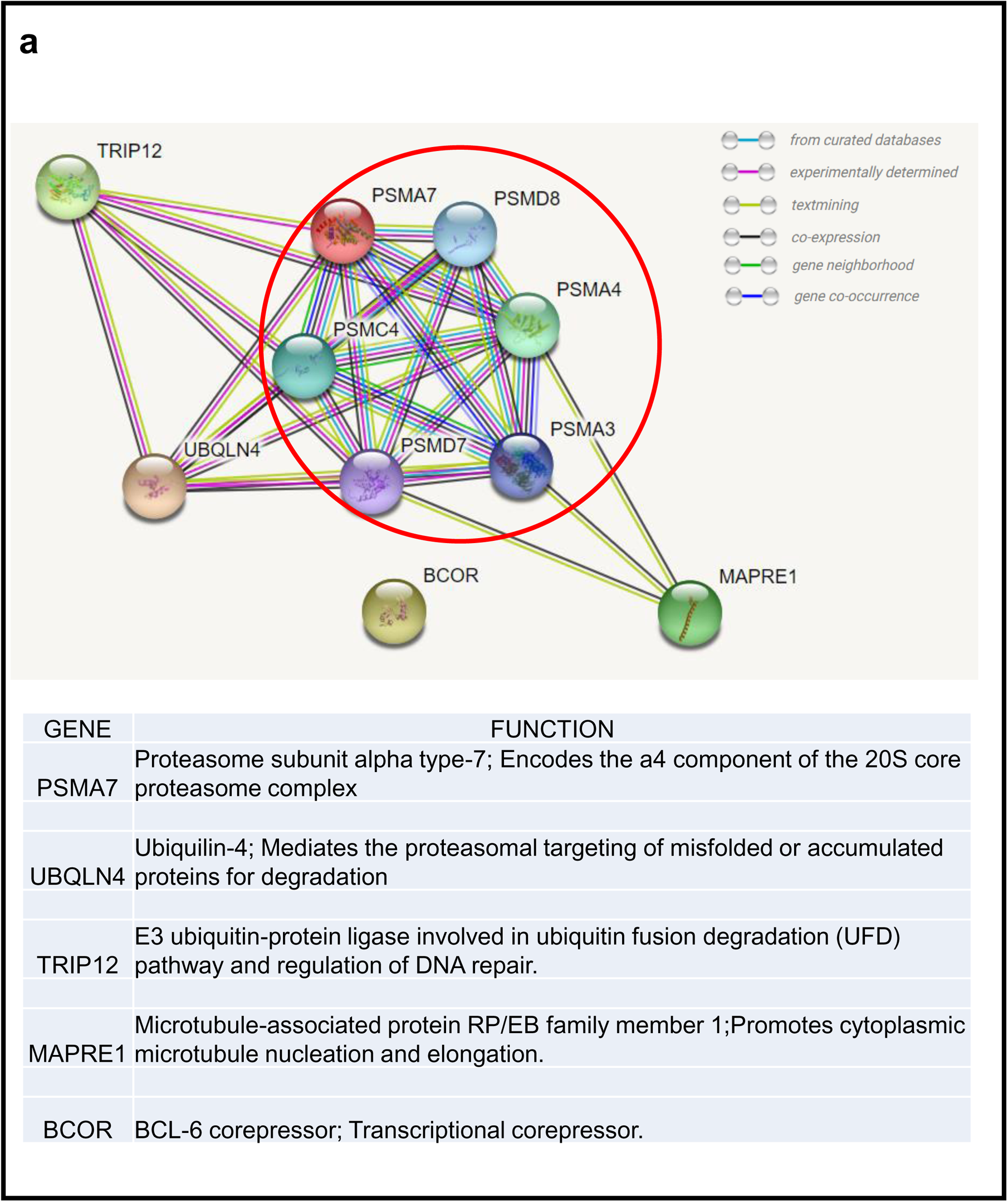
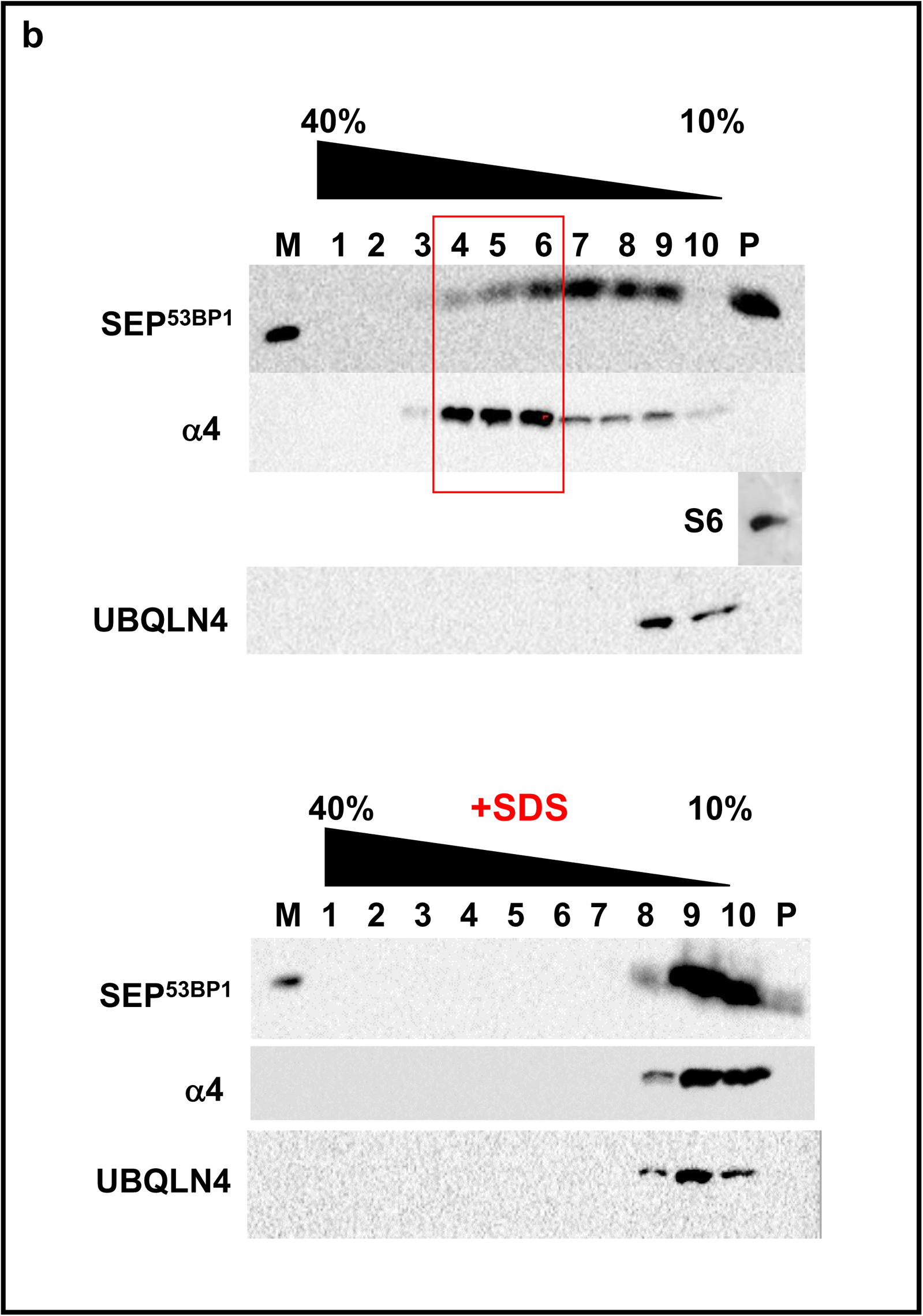
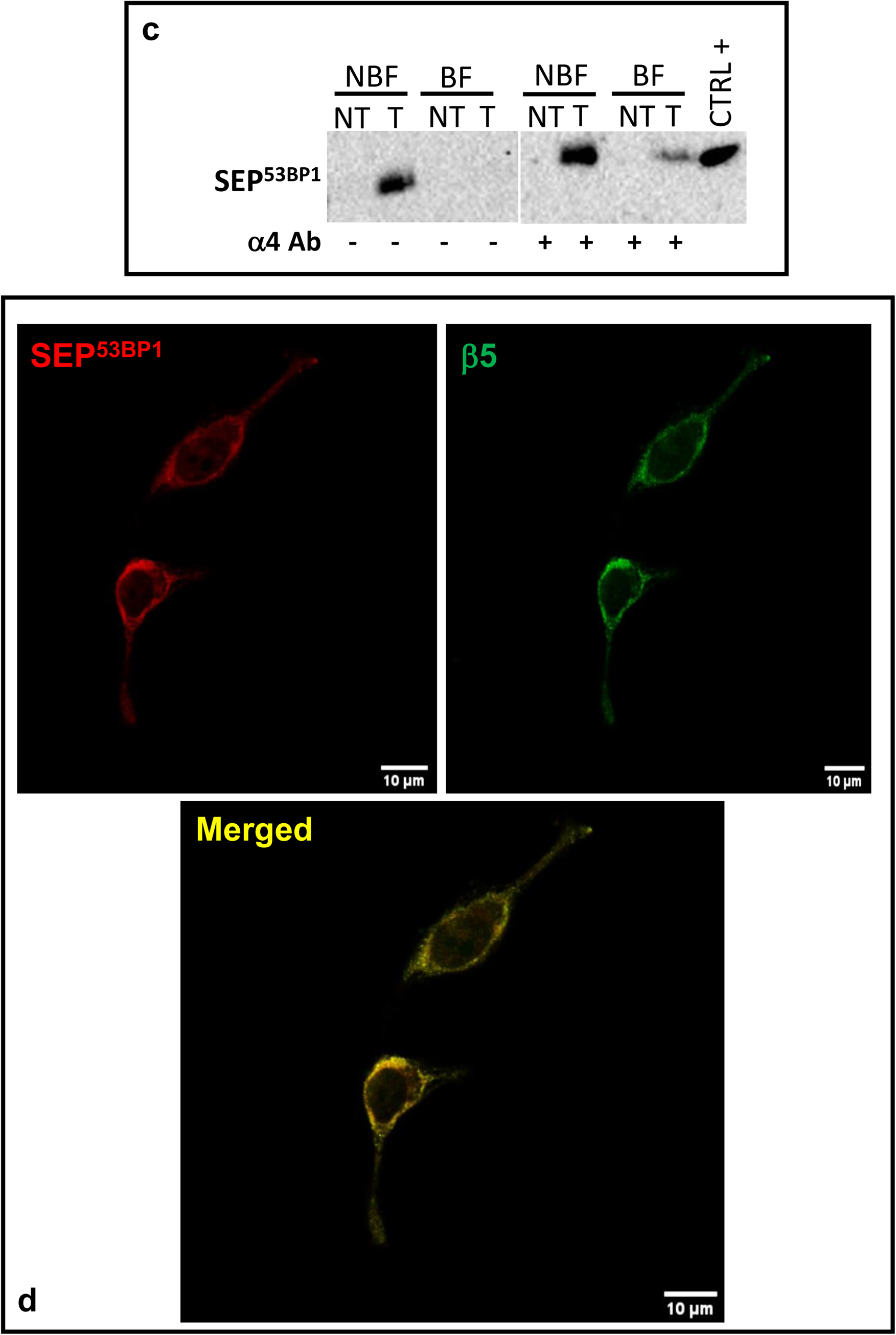
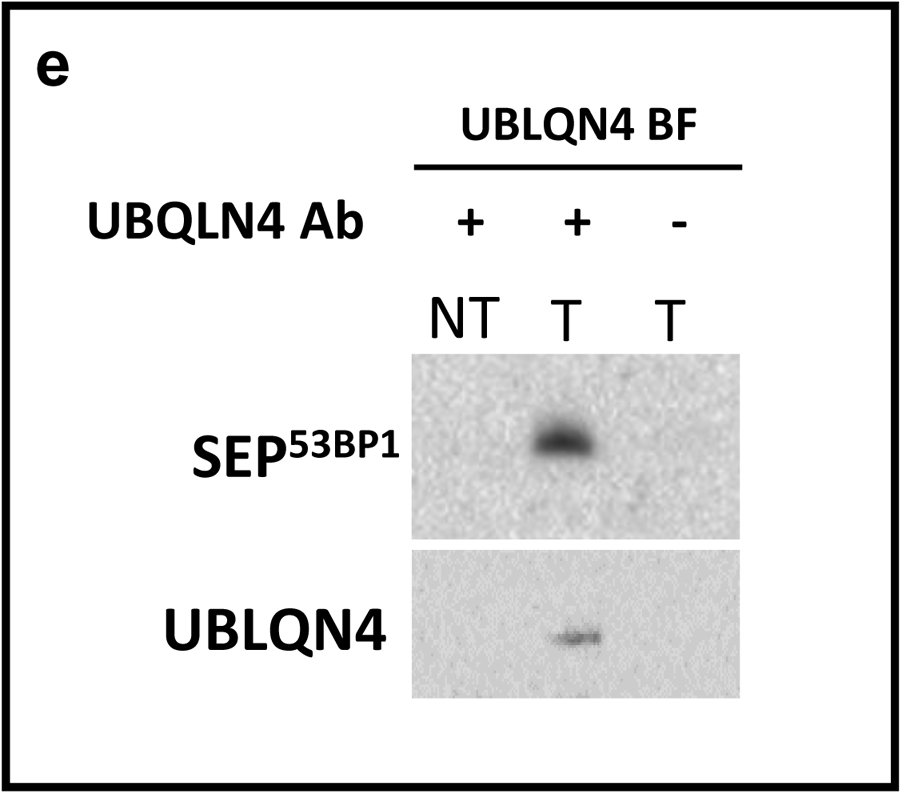
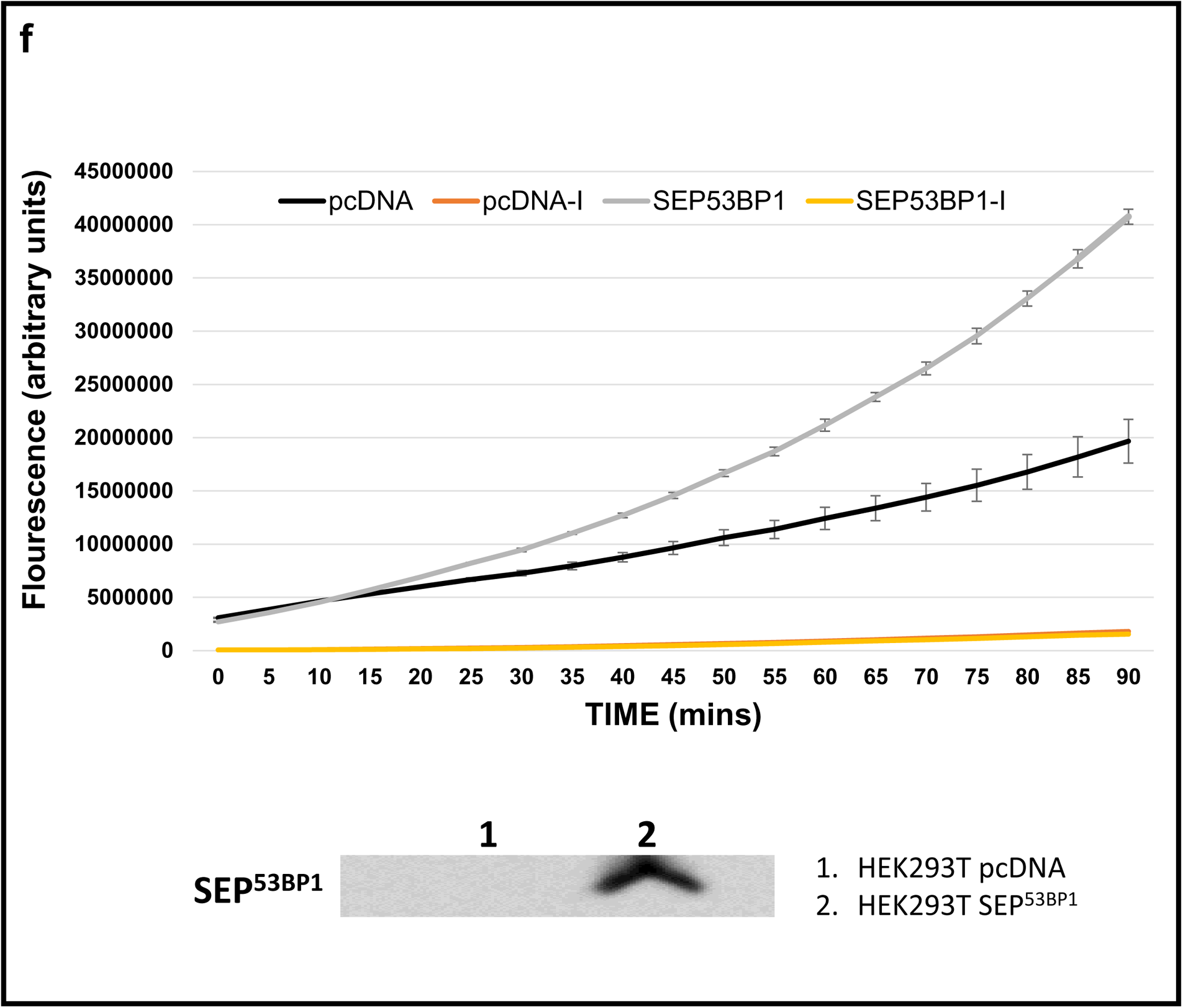
The SEP^53BP1^ interactome. (**a**): String interactions (STRING software) generated using the five SEP^53BP1^ hits from the Y2H analysis as listed in the table. Circled in red are subunits of the proteasome. (**b**) Glycerol gradient profiles generated from HEK293T cells transiently expressing SEP^53BP1^. The cell extracts were prepared in hypotonic lysis buffer supplemented with ATP (see Methods). Ten equal fractions plus the pellet (P) were collected and probed for the α4 and UBQLN4 proteins by Western blotting. The pellet was also analysed for the ribosomal S6 protein. Fractions 4-5-6 are highlighted as the region in which SEP^53BP1^ co-sediments with the majority of the α4 protein. In parallel, lysates were treated with SDS and heated at 65°C prior to loading onto the gradient (lower panels). M indicates a marker for the SEP^53BP1^ protein. (**c**) Immunoblot analysis of the co-IP assay performed on extracts prepared from HEK293T cells transiently expressing SEP^53BP1^ (T) or non-transfected controls (NT). Complexes were recovered on Protein-G magnetic beads alone (-) or beads carrying the α4 antibody (+). NBF and BF indicate the non-binding and binding fractions from the pull-down, respectively. (**d**) HEK293T cells transiently expressing the SEP^53BP1^ protein were grown on glass coverslips. The localisation of SEP^53BP1^ (red staining) and the β5 subunit of the proteasome (green staining) were analysed using confocal IF microscopy using the Zeiss LMS800 confocal scanning microscope. A merged non-contrast-adjusted image is shown in the lower frame. (**e**) Co-IP analysis performed using HEK293T cells transiently expressing both SEP^53BP1^ and UBLQN4 (indicated as T). Non-transfected cells (NT) served as a control. Complexes were recovered on Protein-G magnetic beads alone (-) or beads carrying the UBQLN4 antibody (+) and analysed by immunoblotting with SEP^53BP1^ and UBLQN4 Abs. (**f**) Proteasome activity assay performed using HEK293T lysates prepared from cells transfected with either empty vector (pcDNA) or a vector expressing SEP^53BP1^ (immunoblot presented in the lower panel). The assays indicated with “I” were carried out in the presence of the proteasome inhibitor MG132. All experiments were performed in triplicate and the vertical bars indicate the SEM.

Since the interactome clearly pointed to a role in proteasome biology, we asked if SEP^53BP1^ expression influenced proteasome function. We compared proteasome activity in HEK293T extracts prepared from cells transfected with empty vector or a vector expressing SEP^53BP1^ (Fig. 6f). The presence of the small polypeptide stimulated 26S proteasome activity as confirmed by the inhibitory effect of the drug MG132. The stimulation was greater than two fold over a 90 mins assay period.

## DISCUSSION

It is increasingly evident that the complexity of the metazoan proteome is considerably increased by the expression of SEPs (< 100 aas) that until recently escaped detection using conventional biochemical procedures (they are also referred to as small protein^60^, alt-ORF^61, 62^, nORF^63^, miniproteins or micropeptides^60^). The presence of these products, encoded by smORFs, was initially predicted by Basrai and coworkers^64^, and has subsequently been confirmed using modern techniques such as ribosome profiling, proteogenomics and conservation signatures^60, 65^. The OpenProt database now lists nearly 21,000 human SEPs (www.openprot.org)^66^. They are expressed mainly from lncRNAs and uORFs^24, 67^ but also arise from internal overlapping ORFs or even the 3’UTR, bringing to an end the dogma that eukaryotic mRNAs are monocistronic^25, 38, 39, 42, 68–70^. Furthermore, studies have ascribed diverse functions to these SEPs ranging from the regulation of physiological functions within the cell (e.g. endoplasmic reticulum, Golgi and vesicular transport: see also the introduction)^60, 71^ to crucial developmental functions within metazoan species separated across large evolutionary distances^30, 72–75^.

In this manuscript, we have identified a new member of the SEP family expressed from an ioORF within the 53BP1 gene, the main CDS of which expresses a protein that plays a central role in non-homologous DNA repair^76^. It is the mode of expression and the function of this SEP^53BP1^ that is novel. In humans, it couples alternative promoter activity (P1 versus P2) to a translational reinitiation event on the internal AUG^SEP53BP1^ mediated by a short uORF within the P2 derived mRNA 5’TL. Both these events can respond to intracellular stresses^77–79^. Curiously, it has already been proposed that SEP expression may be an integral part of the cellular “stress response”^80^. The link to developmental functions is also intriguing, because promoter switching and translational reprogramming are key events during differentiation in metazoans ^81, 82^. We have observed P1 promoter activity (V1/2 expression) in all human cell lines tested, indicating that it is the probably the major source of the 53BP1 protein (Fig. 1b). On the other hand, P2 promoter activity showed considerable cell line variability and it remains unclear the molecular basis of its regulation (Fig. 1b). Its V3 transcript directs expression mainly of SEP^53BP1^, indicating that expression of this protein is also not ubiquitous. The low levels that we observed in MCF7 cells (Fig. 5b), despite high cellular levels of the V3 transcript and high polysomal occupancy^47^, suggests that intracellular stability is also regulated in a cell-specific manner. The smORF responsible for SEP expression can be observed in metazoan species from human through to zebrafish (Fig. 3a), with the caveat that part of this conservation may arise from the constraints imposed by the overlapping 53BP1 ORF. However, most of the key functional domains of 53BP1 reside in its C-terminus and there may be more primary sequence plasticity within its long largely disordered N-terminus ^83^. Curiously, zebrafish have only one annotated 53BP1 promoter but the organisation of the 5’TL, more specially the longer uORF, appears to permit ribosome access to both AUG^53BP1^ and AUG^SEP53BP1^ at high efficiency (a sort of fusion of the human V1/2 and V3 readouts: Fig. 3c). This behaviour, with the caveat that our studies have as yet only be performed in mammalian cells, would be consistent with current models of reinitiation^84, 85^ However, the cis-acting sequences on the mRNA that regulate reinitiation, and their responsiveness to intracellular stresses, are well conserved between mammal and zebrafish^8, 86, 87^. Extensive studies on SEP protein expression, and function, have been reported using a *D. melanogaster* model^72, 74, 75, 88^, and riboprofiling studies have confirmed the presence of smORFs in zebrafish^89^. Consequently, zebrafish presents itself as a useful animal model to explore the role of the both uORF and the AUG^SEP53BP1^ in metazoan development.

Our interactome studies revealed that a fraction of the cytoplasmic SEP^53BP1^ associates with the proteasome via its α4 subunit (Fig. 6).The SID on α4 maps to the C-terminus, a region that is found largely exposed on the surface of the 20S and 26S proteasome (Fig.7a/b). This interaction appears to stimulate 26S proteasome activity, based upon the MG132 inhibitory effect^90^, despite the fact that proteolytic activities are on the β rings and positioned in the interior of the 20S cylinder^91^. The 26S specifically targets polyubiquitinylated substrates that are degraded in an ATP-dependent manner (Fig. 6f)^92^. This polyubiquitinylated selectivity resides within the 19S regulatory complex positioned at each extremity of the 20S cylinder (Fig. 7b)^93^. However, during certain stresses, binding of activating proteins can open the α ring on the 20S permitting the entry of protein substrates. Proteins that enter are degraded in an ubiquitin/ATP independent manner. This “active” 20S serves to remove misfolded or oxidised proteins that accumulate during the stress^94^. Furthermore, our detection of SEP^53BP1^ in two cell lines of lymphoid origin is intriguing because lymphoid tissues express a specific proteasome involved in antigen processing called the “immunoproteasome” whose assembly involves changes in the composition of the β ring^95, 96^. It remains to be determined if SEP^53BP1^ is also modulating the activity of the “active” 20S and the immunoproteasome, all of which retain the α4 SID. Proteasomes are found in both cytoplasmic and nuclear compartments^57, 97^. One of the putative nuclear localisation signals is actually located on the C-terminal tail of α4 (Fig. 7a)^98^. It seems conceivable that the nuclear SEP^53BP1^ may enter in association with the proteasome. Furthermore, proteasome levels in the nucleus responds to stresses, such as glucose starvation, hypoxia or low pH,^99^ many of which may also be modulating SEP^53BP1^ intracellular levels.

**Figure 7.**
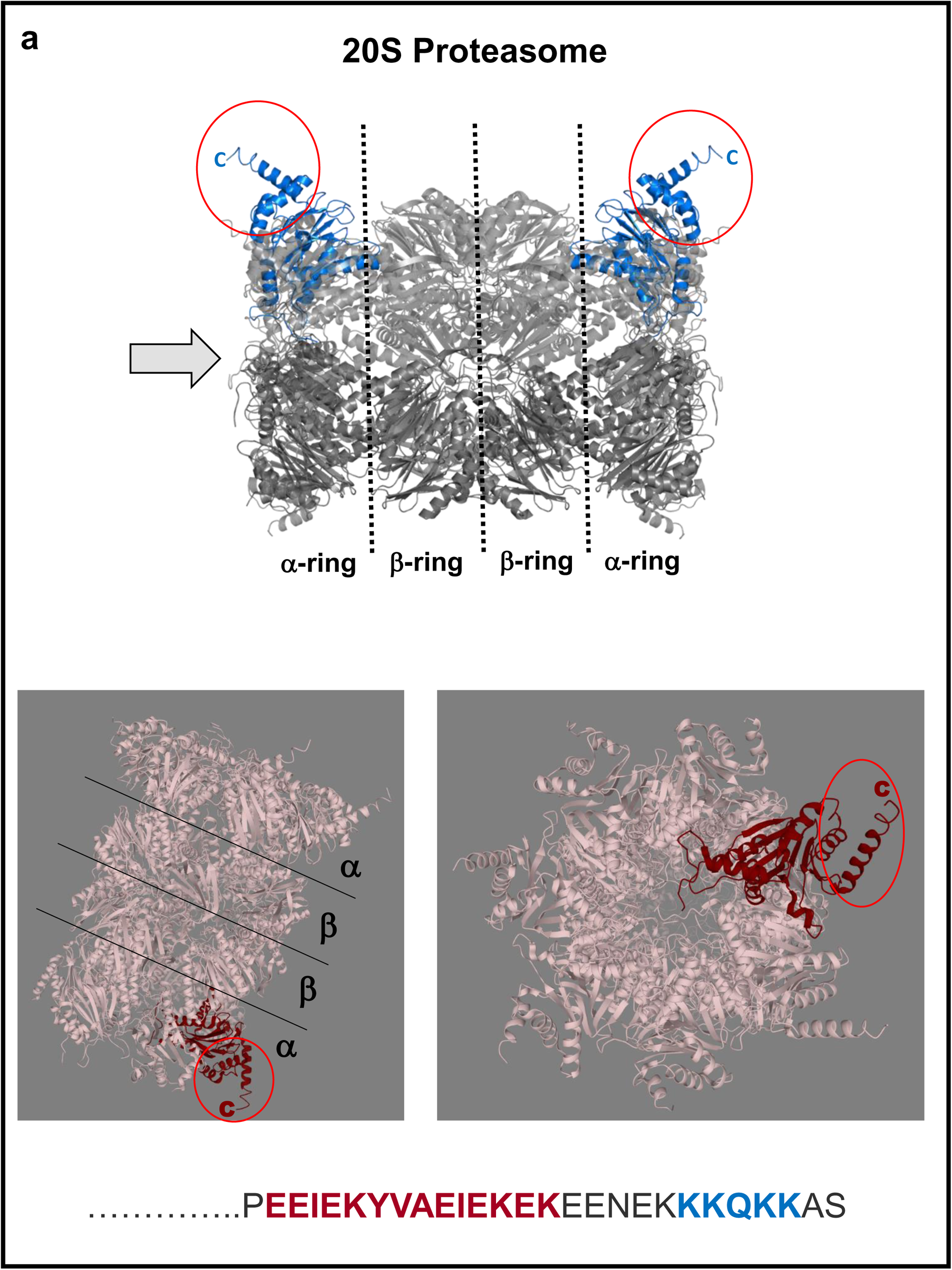
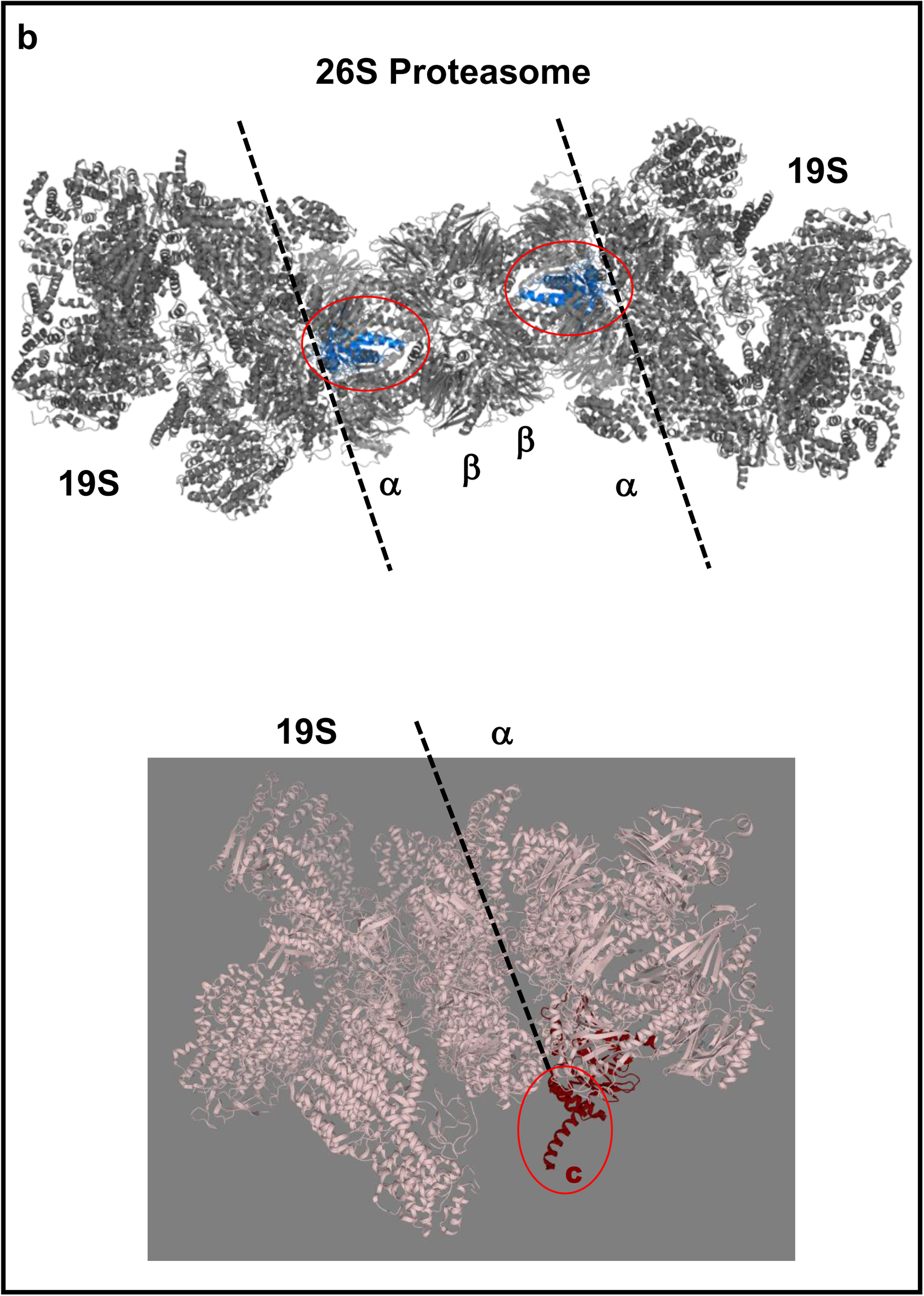
The location of the SEP^53BP1^ interaction site on the human proteasome. (**a**) Upper panel: Side view of the 20S proteasome with the α4 subunits indicated in blue. The exposed C-termini are circled in red. The black dotted lines mark the interface between the α_7_β_7_β_7_α_7_ rings that form the 20S barrel. Lower panels: Rotations of the 20S barrel with α4s indicated in burgundy and the exposed C-termini circled in red. The orientation of the view in the right hand panel (which looks down the 20S barrel) is indicated by the arrow in the upper panel image. The amino acid sequence below is the C-terminal 27 aas of α4. In burgundy is the sequence that forms the extended α-helix visible in all α4 images on the proteasome, and in blue a putative nuclear localisation signal. (**b**) Upper panel: Side view of the 26S proteasome with the α4 subunits indicated in blue. The dotted black line marks the interface between the 20S cylinder and the 19S regulatory caps. Lower panel: Rotation of the 26S showing a 19S-α ring interface (black dotted line) with the exposed C-terminus of α4 circled in red. All images were extracted from the PDBe database (https://www.ebi.ac.uk/pdbe/node/1).

The regulation of proteasome activity has important clinical applications. Proteasome function becomes impaired during many age-related neurodegenerative disorders, including Parkinson’s disease, amyotrophic lateral sclerosis and Alzheimer’s disease. All these conditions are characterised by the accumulation of toxic intracellular protein aggregates that arise because of reduced proteasome activity^100, 101^. Consequently, a considerable amount of research has focused on the development of pharmacological small molecules that can stimulate proteasome function, as potential therapeutic agents for these conditions^48^. However, few of these small molecule compounds activate proteasomes *in vivo*^48^. Increasing P2 promoter activity and intracellular SEP^53BP1^ levels offers a new avenue of research in the treatment of these conditions.

In bicistronic transcripts, the SEP and the product of the CDS frequently exhibit a functionality link. This can involve a direct protein-protein interaction^38, 71, 102, 103^, or alternatively they act indirectly on the same metabolic or physiological pathway^104, 105^. Our Y2H analysis showed no hit with 53BP1 and we have been unable to demonstrate any interaction by co-IP using transiently over-expressed proteins. However, the Y2H study did provide a hit on TRIP12, an E3 ubiquitin-protein ligase that regulates 53BP1 steady state levels in the cell^106^. In a similar vein, UBQLN4 has been reported to co-localise with 53BP1 at the sites of DNA damage and to repress homologous recombination-mediated DNA repair^59^. We have not yet biochemically confirmed these Y2H interactions but it remains possible that SEP^53BP1^ may also impact on the DNA repair process by modulating 53BP1 activity and levels.

### In conclusion

We have identified a new member of the SEP family within the 53BP1 gene, conserved across metazoans. The mode of expression indicates that its intracellular levels will be regulated via alternative promoter activation and cellular stress. One of its functions is to interact with, and modulate, the activity of the cellular proteasome. As such, it offers the possibility of new therapeutic approaches for the treatment of conditions coupled to the accumulation of toxic intracellular protein aggregates.

## METHODS

### Cell culture and transfection

HEK293T, HeLa, MCF7, MCF10A, THP-1 and Raji cells were grown at 37°C in a humidified 5% CO_2_ chamber. Cells were cultured in Dulbecco’s modified Eagle’s medium (DMEM) supplemented with 1% penicillin/streptomycin (P/S) (Gibco) and 10% fœtal bovine serum (FBS, Brunschwig) for HEK293T, HEK293, MDA-MB-231, and HeLa cells. MCF7 (ATCC, HTB-22) were cultured DMEM F-12 medium (Gibco), 1% P/S and 10% FBS supplemented with 10 µg/mL human recombinant insulin and 0.5 nM estradiol. MCF10A (ATCC, CRL-10317) were cultured DMEM F-12 medium containing 1% P/S, 5% horse serum heat inactivated (HS, Brunschwig), 10µg/mL Epidermal Growth Factor (EGF), 1 µM Dexamethasone, and 5µg/ml human recombinant insulin. THP-1 and Raji cells (kind gift of Prof. Walter Reith, UNIGE) were cultured in RPM1 1640 1X (+L-Glutamine) supplemented with 0.05mM 2-mercaptoethanol and 10% fœtal bovine serum. MRC-5 cells were grown in Eagle’s Minimum Essential Medium (MEM; Sigma) supplemented with 10% FBS, 1% P/S, 1X amino acids and 1mM sodium pyruvate. HCT-116 cells were cultured in McCoy’s 5a Medium Modified (Gibco) supplemented with 10% FBS and 1% P/S.

Transfections of HEK293T cells were performed using Lipofectamine 3000 (Invitrogen) when the cells were 70-80% confluent. Eight hours post-transfection, the medium was replaced with normal growth medium, and lysates were usually prepared at 24h post-transfection.

### Drug Treatment

Transfected HEK293T cells were treated for 16 h with DMSO or with 10 µM Cyclopiazonic acid (CPA, Merck) before collection.

### DNA cloning

All clones were prepared in a pcDNA3 backbone. The human α4 20S proteasome subunit was cloned by RT-PCR starting from total cell RNA Trizol extracted from MCF7 cells. The UBQLN4 was subcloned into pcDNA3 starting from the pENTR-A1Up (Plasmid #16170, Addgene). All mutations and deletions were introduced by PCR. The oligos employed are listed in the Supplementary Table 1.

### Polysome gradient/RNA extraction

For polysome profiling, 20–60% sucrose (Sigma) in 100 mM KCl, 5 mM MgCl_2_, 20 mM HEPES and 2 mM DTT gradients were prepared manually in SW41 rotor tubes. Cells were treated for 5 min with 50 μg/ml cycloheximide (Sigma) and then collected in cold PBS containing 100 μg/ml cycloheximide. Cells were pelleted and lysed in polysome lysis buffer (100 mM KCl, 50 mM Tris–HCl pH 7.6, 1.5 mM MgCl_2_, 1 mM DTT, 1 mg/mL Heparin (Sigma), 1.5% (v/v) NP-40, 100 μM cycloheximide, 1% aprotinin, 1 mM AEBSF and 100U/mL of RNasin) supplemented with protease cocktail inhibitor EDTA-free (Roche), on ice for 20 min. Lysates were cleared by centrifugation (14,000xg) and the supernatants loaded onto the gradients. These were centrifuged for three and a half hours at 35,000 rpm and 4°C. They were collected through an UV-lamp and an Absorbance detector. One mL fractions were recovered using a Foxi Junior Fraction Collector (Isco). RNA was isolated from each fraction by adding an equal volume of TriZol (Invitrogen). Samples were mixed and incubated on ice for 15 minutes before addition of 0.3 volumes of chloroform. After centrifugation, the upper phase was collected, and the RNA precipitated with 0.7 volumes of isopropanol. The RNA pellet was resuspended in water.

### RT-PCR

The RT-PCR was performed using 250 ngs of total or polysomal RNA that were reverse transcribed using 50 U of Superscript II (Promega) in a total volume of 25 μL at 42 °C for 1 h. Relative mRNA levels were evaluated by semi-quantitative PCR using the Pfu polymerase (Rovalab) as detailed previously^47^. The number of amplifications cycles was first optimised for each primer set and corresponded to the exponential phase.

### Western blot

Cytoplasmic extracts were prepared in either hypotonic lysis buffer (20mM Hepes, 5mM MgCl_2_, 1mM DTT, 2mM ATP, protease inhibitor) or CSH buffer (50 mM Tris-Cl pH 7.5, 250 mM NaCl, 1 mM EDTA, 0.1% (v/v) Triton X-100). Whole cell extracts were prepared by resuspending the cell pellet in either X2 sample buffer (125 mM Tris-HCl pH 6.8, 20% glycerol, 10% (v/v) β-mercaptoethanol, 5% SDS, 0.025% (w/v) bromophenol blue) for SDS-PAGE, or X2 sample buffer Novex (450mM Tris H-Cl pH 8.45, 12% Glycerol, 4% SDS, 0.00075% Coomassie Blue G, 10% β-mercaptoethanol) for the pre-cast 16% Novex Tricine gels (Invitrogen).

Protein concentrations were determined by Bradford (Cytoskeleton, USA). Twenty μg of protein was resolved on polyacrylamide-SDS gels and electro-transferred to PVDF membranes.

Antibodies used in this study were: anti-53BP1 (Santa Cruz Biotechnologies, sc-22760), anti-phospho (Ser51) eIF2α (GenTex, #61039), anti-eIF2α (Invitrogen, #44728G), anti-HA (Covance clone 16B12), anti-FLAG (M2 antibody, Sigma), anti-actin (Millipore, #MAB1501), anti-RPS6 (Cell Signaling, #2317), anti-A1Up (UBQLN4, Santa Cruz Biotechnologies, sc-136145), anti-α4 20S proteasome (PSMA7, MCP72, Santa Cruz Biotechnologies, sc-58417), and goat anti-mouse or rabbit HRP secondary antibodies (Bio-Rad). The Anti-MYC tag was a gift from Prof. Dominique Soldati (University of Geneva, Switzerland). Immunoblots were developed using the WesternBright™ Quantum (Advansta) and quantitated using Image Lab (Biorad).

### Glycerol gradients

Glycerol gradients were prepared as previously described^107^. HEK293T cells were lysed in hypotonic lysis buffer (see below) containing 2mM ATP. Cell extracts were loaded onto a 10–40% linear glycerol gradient in 100 mM KCl, 5 mM MgCl_2_, and 20 mM HEPES pH 7.4 prepared in an SW60 tube. Gradients were centrifuged for 16 hr at 30,000 rpm in a SW60 rotor at room temperature. After centrifugation, 10 X 400 μL fractions were collected from the bottom of the tube. Proteins were recovered by methanol/chloroform precipitation, resuspended directly in 40μL of X2 sample buffer and analysed by western blotting.

### Preparation of hypotonic cell extracts

Extracts were prepared following the protocol outlined in Terenin and coworkers^108^. Actively dividing cells (≈70% confluence) from a 100mm petri dishe were scrapped into ice-cold PBS (final volume 1 mL) and recovered by pelleting at 1,000g for 5 mins at 4°C. They were washed a second time with PBS before resuspending in 200 μL of Lysolecitin lysis buffer (20 mM HEPES-KOH pH 7.4, 100 mM KOAc, 20 mM DTT, 0.1 mg/mL (w/v) Lysolecitin) for precisely 1 minute on ice before centrifuging at 10,000g for 10 secs at 4°C. The pellet was resuspended in an equal volume of ice-cold hypotonic extraction buffer (20 mM HEPES-KOH pH 7.5, 10 mM KOAc, 1 mM MgAc, 4 mM DTT containing complete protease inhibitor cocktail-EDTA minus). Cells were disrupted in a pre-cooled Dounce homogeniser using 20-25 strokes, transferred to an Eppendorf tube, and spun at 10,000g for 10 mins at 4°C.

### Yeast-two-hybrid

Y2H was performed by Hybrigenics Services (https://www.hybrigenics-services.com/) against the Human B cell Lymphoma library.

### Co-immunoprecipitation

*α4 pull-down:* SEP^53BP1^ was transiently expressed in HEK293T cells. Cells were lysed in hypotonic lysis buffer containing 2mM ATP. 500 μg of protein were pre-incubated with α4 antibody overnight at 4°C. The next day, the lysates were incubated with 10 μL of Dynabeads Protein G (Invitrogen) for 1 hr at 4°C. Beads were washed X3 in hypotonic lysis buffer, resuspended in X2 sample buffer and analysed by SDS-PAGE.

*UBQLN4 pull-down:* HEK293T cells were co-transfected with SEP^53BP1^ and UBQLN4. They were lysed in hypotonic lysis buffer. 500 μg of protein were pre-incubated with UBQLN4 antibody overnight at 4°C. The next day, the lysates were incubated with 10 μL of Dynabeads Protein G (Invitrogen) 1 hr at 4°C. Beads were washed X3 in hypotonic lysis buffer, resuspended in X2 sample buffer and analysed by SDS-PAGE.

### *In-vitro* transcription and translation

Plasmids were linearized and RNA *in vitro* transcribed using the mMessage mMachine™ T7 Transcription Kit (Invitrogen) and purified by LiCl precipitation according to the manufacturer’s instructions. Transcripts were capped using the vaccinia capping system (NEB) according to the suppliers’ protocol and polyadenylated using the polyA tailing kit (Invitrogen). The mRNA quality was controlled on a 2% agarose gel. RNA (100 ng) was translated in a 50% wheat germ extract (WGE, Promega) for 90 mins at 25°C in a final volume of 20 μL. The reaction was stopped by the addition of Laemmli buffer SB (X4 buffer: 250 mM Tris HCl pH 6.8, 40% glycerol, 20% β-mercaptoethanol, 10% SDS, 0.05% Bromophenol blue.

### Indirect immunofluorescence using confocal imaging and z-stacking

SEP^53BP1^ was transiently expressed in HEK293T cells grown on No. 1.5 coverslips. Cell suspensions of THP-1 and Raji were pelleted and resuspended in PBS. They were allowed to adhere on the coverslips by sedimentation for 30 minutes at RT. Cells were fixed with 4% PFA, permeabilised with 0.3% Triton/PBS and then incubated in blocking buffer (2% BSA/PBS) for 60 minutes at RT. Blocking buffer was replaced by the SEP^53BP1^ primary antibody (1:100 dilution in blocking buffer) overnight at 4°C. Cells were washed X3 with PBS and incubated in the dark for 60 minutes at RT with Alexa Fluor 594 goat anti-rabbit IgG (H+L) (1:500 dilution in blocking buffer) (Invitrogen). Coverslips were washed X3 with PBS. Nuclei were stained with DAPI (1:1000 in PBS) at RT for 10 minutes in the dark. Coverslips were mounted using Prolong Diamond Antifade Mountant (Invitrogen) and sealed with nail polish.

Confocal images were collected on a Zeiss LSM800 confocal scanning microscope equipped with a Plan-Apochromat 63x/1.40 Oil DIC M27 objective. Pictures were analysed using the ImageJ software.

Z-series videos in THP1 and RAJI cells are shown as maximum z-projections, and gamma, brightness, and contrast were adjusted (identically for compared image sets) and were generated using the Imaris software.

### Proteasomal activity assay

The proteasomal activity assay was performed on HEK293T cells transfected with either empty vector or a vector expressing SEP^53BP1^. Activity was measured using the Proteasome Activity kit (ab107921, Abcam) following the manufacturer’s protocol. MG-132 (Sigma) treatment served as a control to differentiate between 26S proteasome activity and other protease activities present in the extract. The reaction was followed for 90 minutes, and fluorescence was measured on a fluorometric microplate reader (SpectraMax Paradigm) equipped with an Ex/Em: 350/440nm filter, R.E.A.D.S. Unit Experiments were performed in triplicate.

## Supporting information

Supplemental animation 1

Supplemental animation 2

## Acknowledgements

This work was supported by the University of Geneva, the Swiss Science Foundation (31003A_175560) the Société Académique de Genève, the Ernst and Lucie Schmidheiny Foundation, the Ligue Genevoise Contre le Cancer and the Fondation Pour la Lutte Contre le Cancer. We would like to thank Dr. Richard Fish (UNIGE) for supplying us with total zebrafish RNA and Prof. Dominique Belin for helpful discussions.

## Author contributions

M.A.I., M.A., T.H and P.J-G performed the experiments. J.A.C designed the project, interpreted the data and wrote the article. J.A.C. supervised the project.

## Additional information

Supplementary Information accompanies this paper.

## Supplementary Figure Legends

**Supplementary Figure 1:**
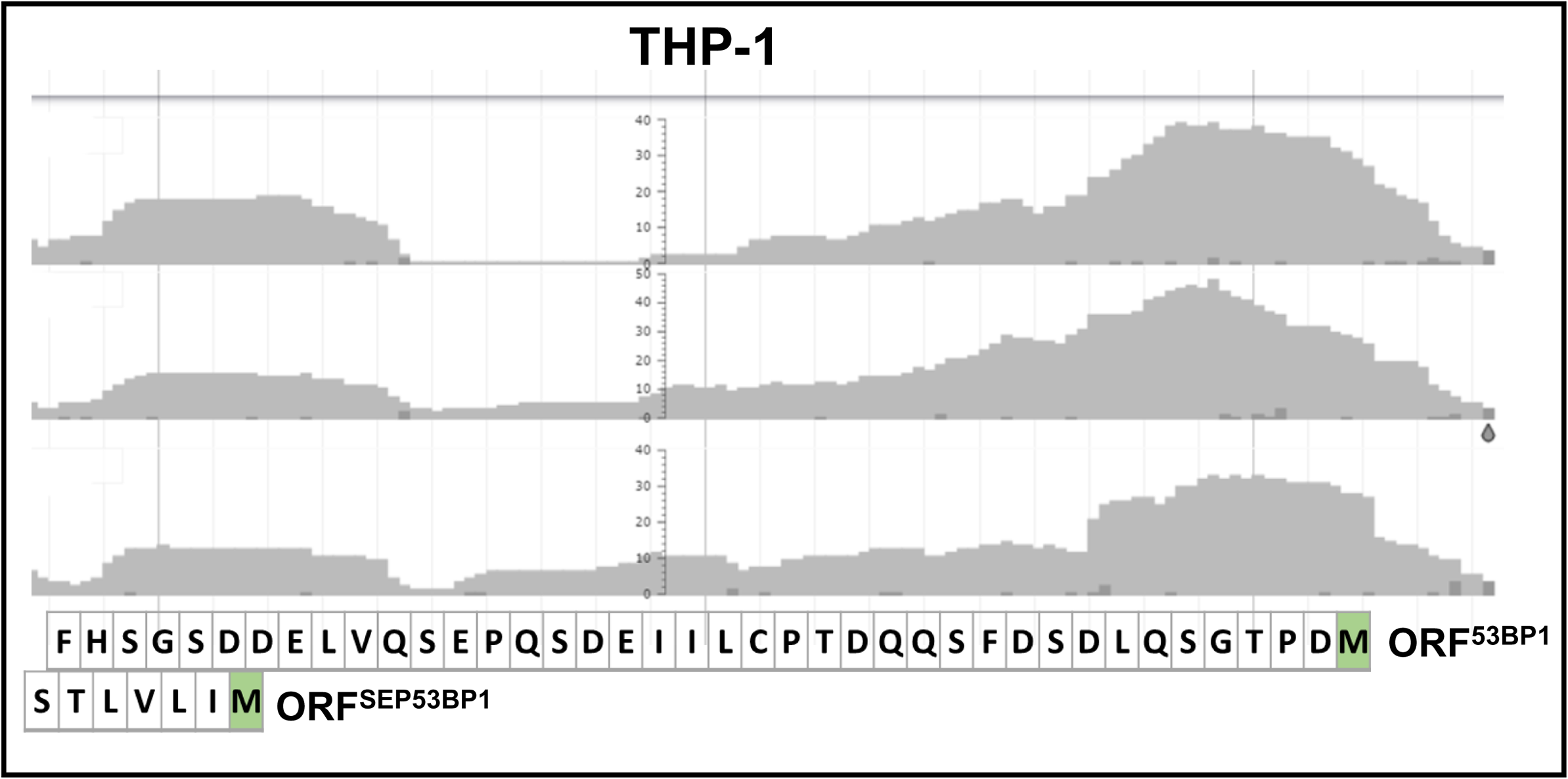
Ribosome Profiling analysis of 53BP1 gene expression in THP-1 cells. The data was extracted from the RPF database http://sysbio.sysu.edu.cn/rpfdb/index.html. The upper graph shows read density mapped through the AUG^53BP1^ and AUG^SEP53BP1^ start sites from three independent experiments. The mRNA is decoded from left to right as indicated by the corresponding amino acid sequence for the two overlapping reading frames. The N-terminal methionine’s for 53BP1 and SEP53BP1 are highlighted in green.

**Supplementary Figure 2:**
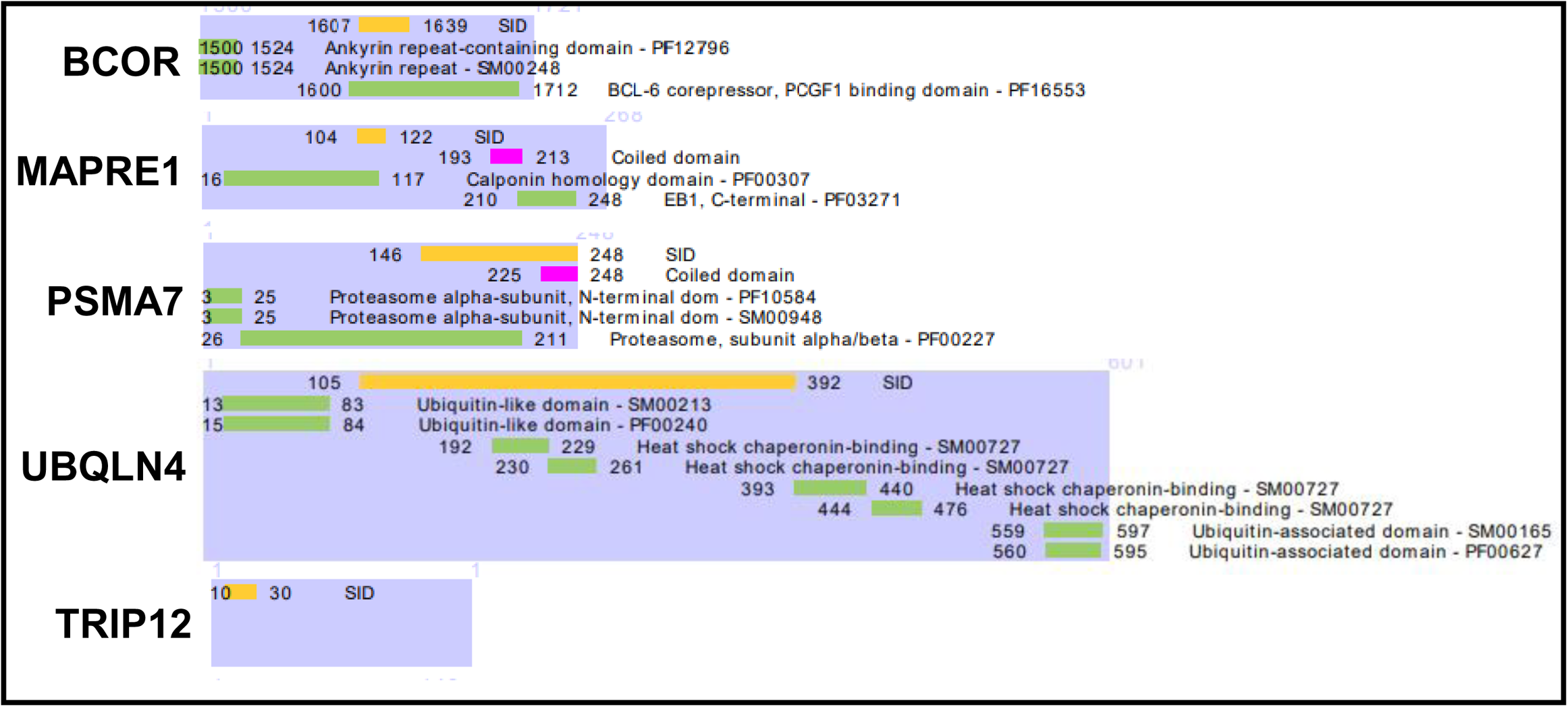
The SEP^53BP1^ partners as determined by Y2H are represented according to their protein chain length. The selected interaction domains (SID) and the known functional domains of each protein are all indicated.

**Supplementary animation 1**: Cellular distribution of endogenous SEP^53BP1^ in THP-1 cells.

**Supplementary animation 2**: Cellular distribution of endogenous SEP^53BP1^ in Raji cells.

Z-stacking was performed using a Zeiss LSM800 confocal scanning microscope equipped with a Plan-Apochromat 63x/1.40 Oil DIC M27 objective. For THP-1 (Supplementary animation 1) 38 slices were taken (7.77 µm) and for Raji (Supplementary animation 2) 30 slices (6,09 µm), in a manner that assures that the cell is imaged in its entirety. Pictures were analysed by the Imaris software. The first few seconds of each video shows the original pictures collected, with the endogenous SEP^53BP1^ staining red and the nuclei staining blue.

This is followed by a 3-Dimensional reconstruction of the nucleus using the surface function within Imaris. In this representation, the cytoplasmic SEP^53BP1^ stains magenta and nuclear SEP^53BP1^ stains light blue.

**Supplementary table 1:**
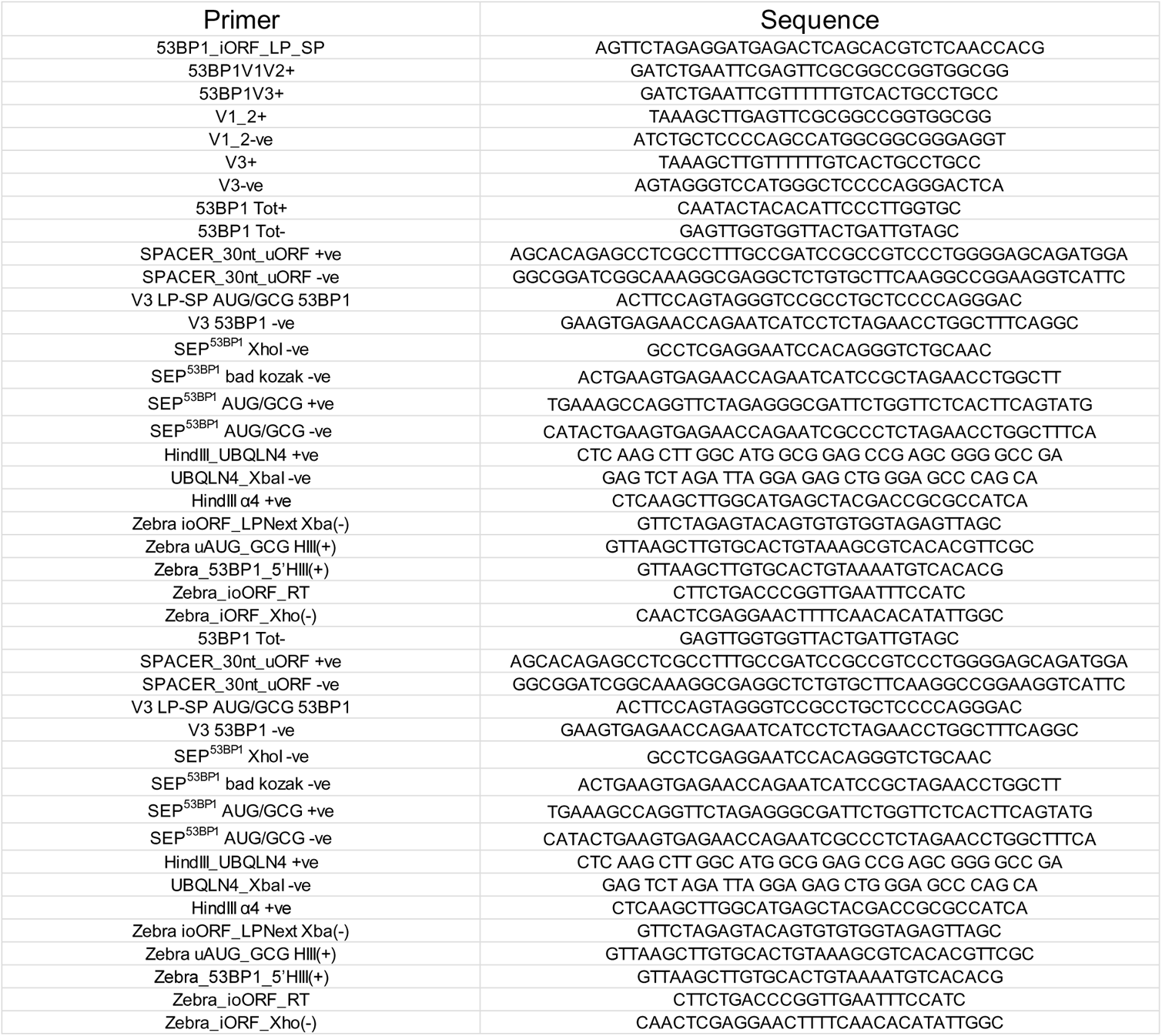
List of oligos used.

## REFERENCES

1. Ingolia, Nicholas T., Lareau, Liana F. & Weissman, Jonathan S. Ribosome Profiling of Mouse Embryonic Stem Cells Reveals the Complexity and Dynamics of Mammalian Proteomes. Cell 147, 789–802, doi:10.1016/j.cell.2011.10.002 (2011).

2. Kozak, M. Point mutations define a sequence flanking the AUG initiator codon that modulates translation by eukaryotic ribosomes. Cell 44, 283–292 (1986).

3. Wang, X. Q. & Rothnagel, J. A. 5’-Untranslated regions with multiple upstream AUG codons can support low-level translation via leaky scanning and reinitiation. Nucleic Acids Research 32, 1382–1391 (2004).

4. Curran, J. A., Richardson, C. & Kolakofsky, D. Ribosomal initiation at alternate AUGs on the Sendai virus P/C mRNA. J.Virol. 57, 684–687 (1986).

5. Hinnebusch, A. G., Ivanov, I. P. & Sonenberg, N. Translational control by 5′-untranslated regions of eukaryotic mRNAs. Science 352, 1413–1416, doi:10.1126/science.aad9868 (2016).

6. Ingolia, N. T., Ghaemmaghami, S., Newman, J. R. & Weissman, J. S. Genome-wide analysis in vivo of translation with nucleotide resolution using ribosome profiling. Science 324, doi:10.1126/science.1168978 (2009).

7. Andreev, D. E. et al. Translation of 5’ leaders is pervasive in genes resistant to eIF2 repression. Elife 4, e03971, doi:10.7554/eLife.03971 (2015).

8. Johnstone, T. G., Bazzini, A. A. & Giraldez, A. J. Upstream ORFs are prevalent translational repressors in vertebrates. 35, 706–723, doi:10.15252/embj.201592759 (2016).

9. Morris, D. R. & Geballe, A. P. Upstream open reading frames as regulators of mRNA translation. Mol.Cell Biol. 20, 8635–8642 (2000).

10. Rahim, G., Araud, T., Jaquier-Gubler, P. & Curran, J. Alternative splicing within the elk-1 5’ untranslated region serves to modulate initiation events downstream of the highly conserved upstream open reading frame 2. Mol Cell Biol 32, 1745–1756, doi:10.1128/MCB.06751-11 (2012).

11. Chew, G.-L., Pauli, A. & Schier, A. F. Conservation of uORF repressiveness and sequence features in mouse, human and zebrafish. Nature Communications 7, 11663, doi:10.1038/ncomms11663 (2016).

12. Proud, C. G. eIF2 and the control of cell physiology. Semin.Cell Dev Biol 16, 3–12, doi:S1084-9521(04)00106-5 [pii];10.1016/j.semcdb.2004.11.004 [doi] (2005).

13. Hinnebusch, A. G. The eIF-2 alpha kinases: regulators of protein synthesis in starvation and stress. Semin.Cell Biol. 5, 417–426 (1994).

14. Pal, S. et al. Alternative transcription exceeds alternative splicing in generating the transcriptome diversity of cerebellar development. 21, 1260–1272, doi:10.1101/gr.120535.111 (2011).

15. Stamatoyannopoulos, J. A. Illuminating eukaryotic transcription start sites. Nature Methods 7, 501, doi:10.1038/nmeth0710-501 (2010).

16. Curran, J. A. & Weiss, B. What Is the Impact of mRNA 5’ TL Heterogeneity on Translational Start Site Selection and the Mammalian Cellular Phenotype? Front Genet 7, 156, doi:10.3389/fgene.2016.00156 (2016).

17. Landry, J.-R., Mager, D. L. & Wilhelm, B. T. Complex controls: the role of alternative promoters in mammalian genomes. Trends in Genetics 19, 640–648, doi:10.1016/j.tig.2003.09.014 (2003).

18. Calvo, S. E., Pagliarini, D. J. & Mootha, V. K. Upstream open reading frames cause widespread reduction of protein expression and are polymorphic among humans. Proc.Natl.Acad.Sci.U.S.A 106, 7507–7512, doi:0810916106 [pii];10.1073/pnas.0810916106 [doi] (2009).

19. Somers, J., Pöyry, T. & Willis, A. E. A perspective on mammalian upstream open reading frame function. Int J Biochem Cell Biol 45, 1690–1700, doi:10.1016/j.biocel.2013.04.020 (2013).

20. Brown, C. Y., Mize, G. J., Pineda, M., George, D. L. & Morris, D. R. Role of two upstream open reading frames in the translational control of oncogene mdm2. Oncogene 18, 5631–5637 (1999).

21. Genolet, R., Rahim, G., Gubler-Jaquier, P. & Curran, J. The translational response of the human mdm2 gene in HEK293T cells exposed to rapamycin: a role for the 5’-UTRs. Nucleic Acids Res 39, 989–1003, doi:10.1093/nar/gkq805 (2011).

22. Rajasekhar, V. K. et al. Oncogenic Ras and Akt signaling contribute to glioblastoma formation by differential recruitment of existing mRNAs to polysomes. Mol.Cell 12, 889–901 (2003).

23. Saghatelian, A. & Couso, J. P. Discovery and characterization of smORF-encoded bioactive polypeptides. Nature chemical biology 11, 909–916, doi:10.1038/nchembio.1964 (2015).

24. Slavoff, S. A. et al. Peptidomic discovery of short open reading frame-encoded peptides in human cells. Nature chemical biology 9, 59–64, doi:10.1038/nchembio.1120 (2013).

25. Vanderperre, B. et al. Direct detection of alternative open reading frames translation products in human significantly expands the proteome. PLoS One 8, e70698, doi:10.1371/journal.pone.0070698 (2013).

26. Martinez, T. F. et al. Accurate annotation of human protein-coding small open reading frames. Nature Chemical Biology 16, 458–468, doi:10.1038/s41589-019-0425-0 (2020).

27. Couso, J. P. Finding smORFs: getting closer. Genome Biology 16, 189, doi:10.1186/s13059-015-0765-3 (2015).

28. Mumtaz, M. Ali S. & Couso, Juan P. Ribosomal profiling adds new coding sequences to the proteome. Biochemical Society Transactions 43, 1271–1276, doi:10.1042/BST20150170 %J Biochemical Society Transactions (2015).

29. Ji, Z., Song, R., Regev, A. & Struhl, K. Many lncRNAs, 5’UTRs, and pseudogenes are translated and some are likely to express functional proteins. eLife 4, e08890, doi:10.7554/eLife.08890 (2015).

30. Andrews, S. J. & Rothnagel, J. A. Emerging evidence for functional peptides encoded by short open reading frames. Nat Rev Genet 15, 193–204, doi:10.1038/nrg3520 (2014).

31. Ivanov, I. P. et al. Polyamine Control of Translation Elongation Regulates Start Site Selection on Antizyme Inhibitor mRNA via Ribosome Queuing. Molecular Cell 70, 254–264.e256, doi:https://doi.org/10.1016/j.molcel.2018.03.015 (2018).

32. Ivanova, I. G., Park, C. V., Yemm, A. I. & Kenneth, N. S. PERK/eIF2α signaling inhibits HIF-induced gene expression during the unfolded protein response via YB1-dependent regulation of HIF1α translation. Nucleic Acids Research, gky127–gky127, doi:10.1093/nar/gky127 (2018).

33. Jorgensen, R. A. & Dorantes-Acosta, A. E. Conserved Peptide Upstream Open Reading Frames are Associated with Regulatory Genes in Angiosperms. Frontiers in plant science 3, 191, doi:10.3389/fpls.2012.00191 (2012).

34. Ivanov, I. P., Firth, A. E., Michel, A. M., Atkins, J. F. & Baranov, P. V. Identification of evolutionarily conserved non-AUG-initiated N-terminal extensions in human coding sequences. Nucleic Acids Res 39, 4220–4234, doi:10.1093/nar/gkr007 (2011).

35. Van Damme, P., Gawron, D., Van Criekinge, W. & Menschaert, G. N-terminal Proteomics and Ribosome Profiling Provide a Comprehensive View of the Alternative Translation Initiation Landscape in Mice and Men. Molecular &amp;amp; Cellular Proteomics 13, 1245, doi:10.1074/mcp.M113.036442 (2014).

36. Giorgi, C., Blumberg, B. M. & Kolakofsky, D. Sendai virus contains overlapping genes expressed from a single mRNA. Cell 35, 829–836 (1983).

37. Mouilleron, H., Delcourt, V. & Roucou, X. Death of a dogma: eukaryotic mRNAs can code for more than one protein. Nucleic Acids Res 44, 14–23, doi:10.1093/nar/gkv1218 (2016).

38. Bergeron, D. et al. An out-of-frame overlapping reading frame in the ataxin-1 coding sequence encodes a novel ataxin-1 interacting protein. J Biol Chem 288, 21824–21835, doi:10.1074/jbc.M113.472654 (2013).

39. Vanderperre, B. et al. An overlapping reading frame in the PRNP gene encodes a novel polypeptide distinct from the prion protein. 25, 2373–2386, doi:10.1096/fj.10-173815 (2011).

40. Payre, F. & Desplan, C. RNA. Small peptides control heart activity. Science (New York, N.Y.) 351, 226–227, doi:10.1126/science.aad9873 (2016).

41. Guo, B. et al. Humanin peptide suppresses apoptosis by interfering with Bax activation. Nature 423, 456–461, doi:10.1038/nature01627 (2003).

42. Slavoff, S. A., Heo, J., Budnik, B. A., Hanakahi, L. A. & Saghatelian, A. A human short open reading frame (sORF)-encoded polypeptide that stimulates DNA end joining. J Biol Chem 289, 10950–10957, doi:10.1074/jbc.C113.533968 (2014).

43. Khitun, A., Ness, T. J. & Slavoff, S. A. Small open reading frames and cellular stress responses. Molecular omics 15, 108–116, doi:10.1039/c8mo00283e (2019).

44. Oh, S. et al. Human CTLs to Wild-Type and Enhanced Epitopes of a Novel Prostate and Breast Tumor-Associated Protein, TARP, Lyse Human Breast Cancer Cells. Cancer Research 64, 2610, doi:10.1158/0008-5472.CAN-03-2183 (2004).

45. Slager, E. H. et al. CD4&lt;sup&gt;+&lt;/sup&gt; Th2 Cell Recognition of HLA-DR-Restricted Epitopes Derived from CAMEL: A Tumor Antigen Translated in an Alternative Open Reading Frame. The Journal of Immunology 170, 1490, doi:10.4049/jimmunol.170.3.1490 (2003).

46. Sendoel, A. et al. Translation from unconventional 5′ start sites drives tumour initiation. Nature 541, 494, doi:10.1038/nature21036 https://www.nature.com/articles/nature21036#supplementary-information (2017).

47. Dieudonne, F. X. et al. The effect of heterogeneous Transcription Start Sites (TSS) on the translatome: implications for the mammalian cellular phenotype. BMC Genomics 16, 986, doi:10.1186/s12864-015-2179-8 (2015).

48. Leestemaker, Y. et al. Proteasome Activation by Small Molecules. Cell Chemical Biology 24, 725–736.e727, doi:https://doi.org/10.1016/j.chembiol.2017.05.010 (2017).

49. Wiesenthal, V., Leutz, A. & Calkhoven, C. F. Analysis of translation initiation using a translation control reporter system. Nat.Protoc. 1, 1531–1537, doi:nprot.2006.274 [pii];10.1038/nprot.2006.274 [doi] (2006).

50. Guan, B.-J. et al. A Unique ISR Program Determines Cellular Responses to Chronic Stress. Molecular Cell 68, 885–900.e886, doi:https://doi.org/10.1016/j.molcel.2017.11.007 (2017).

51. Fritsch, C. et al. Genome-wide search for novel human uORFs and N-terminal protein extensions using ribosomal footprinting. Genome Res 22, 2208–2218, doi:10.1101/gr.139568.112 (2012).

52. Wolin, S. L. & Walter, P. Ribosome pausing and stacking during translation of a eukaryotic mRNA. Embo j 7, 3559–3569 (1988).

53. Jackson, R. & Standart, N. The awesome power of ribosome profiling. 21, 652–654, doi:10.1261/rna.049908.115 (2015).

54. Apcher, G.-S., Maitland, J., Dawson, S., Sheppard, P. & Mayer, R. J. The α4 and α7 subunits and assembly of the 20S proteasome. FEBS Letters 569, 211–216, doi:https://doi.org/10.1016/j.febslet.2004.05.067 (2004).

55. Hirano, Y. et al. A heterodimeric complex that promotes the assembly of mammalian 20S proteasomes. Nature 437, 1381–1385, doi:10.1038/nature04106 (2005).

56. Orino, E. et al. ATP-dependent reversible association of proteasomes with multiple protein components to form 26S complexes that degrade ubiquitinated proteins in human HL-60 cells. FEBS Lett 284, 206–210, doi:10.1016/0014-5793(91)80686-w (1991).

57. Peters, J. M., Franke, W. W. & Kleinschmidt, J. A. Distinct 19 S and 20 S subcomplexes of the 26 S proteasome and their distribution in the nucleus and the cytoplasm. J Biol Chem 269, 7709–7718 (1994).

58. Hirayama, S. et al. Nuclear export of ubiquitinated proteins via the UBIN-POST system. Proc Natl Acad Sci U S A 115, E4199–e4208, doi:10.1073/pnas.1711017115 (2018).

59. Jachimowicz, R. D. et al. UBQLN4 Represses Homologous Recombination and Is Overexpressed in Aggressive Tumors. Cell 176, 505–519.e522, doi:10.1016/j.cell.2018.11.024 (2019).

60. Orr, M. W., Mao, Y., Storz, G. & Qian, S. B. Alternative ORFs and small ORFs: shedding light on the dark proteome. Nucleic Acids Res 48, 1029–1042, doi:10.1093/nar/gkz734 (2020).

61. Vanderperre, B. et al. Direct Detection of Alternative Open Reading Frames Translation Products in Human Significantly Expands the Proteome. PLOS ONE 8, e70698, doi:10.1371/journal.pone.0070698 (2013).

62. Menschaert, G. et al. Deep proteome coverage based on ribosome profiling aids mass spectrometry-based protein and peptide discovery and provides evidence of alternative translation products and near-cognate translation initiation events. Mol Cell Proteomics 12, 1780–1790, doi:10.1074/mcp.M113.027540 (2013).

63. Neville, M. D. C. et al. A platform for curated products from novel open reading frames prompts reinterpretation of disease variants. Genome Res, doi:10.1101/gr.263202.120 (2021).

64. Basrai, M. A., Hieter, P. & Boeke, J. D. Small open reading frames: beautiful needles in the haystack. Genome Res 7, 768–771, doi:10.1101/gr.7.8.768 (1997).

65. Pueyo, J. I., Magny, E. G. & Couso, J. P. New Peptides Under the s(ORF)ace of the Genome. Trends Biochem Sci 41, 665–678, doi:10.1016/j.tibs.2016.05.003 (2016).

66. Brunet, M. A. et al. OpenProt: a more comprehensive guide to explore eukaryotic coding potential and proteomes. Nucleic Acids Res 47, D403–D410, doi:10.1093/nar/gky936 (2019).

67. Oyama, M. et al. Analysis of Small Human Proteins Reveals the Translation of Upstream Open Reading Frames of mRNAs. Genome Research 14, 2048–2052 (2004).

68. Beadle, G. W. & Tatum, E. L. Genetic Control of Biochemical Reactions in Neurospora. Proc Natl Acad Sci U S A 27, 499–506, doi:10.1073/pnas.27.11.499 (1941).

69. Brunet, M. A., Levesque, S. A., Hunting, D. J., Cohen, A. A. & Roucou, X. Recognition of the polycistronic nature of human genes is critical to understanding the genotype-phenotype relationship. Genome research 28, 609–624, doi:10.1101/gr.230938.117 (2018).

70. Ingolia, N. T. Ribosome Footprint Profiling of Translation throughout the Genome. Cell 165, 22–33, doi:10.1016/j.cell.2016.02.066 (2016).

71. Chen, J. et al. Pervasive functional translation of noncanonical human open reading frames. Science 367, 1140–1146, doi:10.1126/science.aay0262 (2020).

72. Galindo, M. I., Pueyo, J. I., Fouix, S., Bishop, S. A. & Couso, J. P. Peptides encoded by short ORFs control development and define a new eukaryotic gene family. PLoS Biol 5, e106, doi:10.1371/journal.pbio.0050106 (2007).

73. Magny, E. G. et al. Conserved Regulation of Cardiac Calcium Uptake by Peptides Encoded in Small Open Reading Frames. Science 341, 1116, doi:10.1126/science.1238802 (2013).

74. Couso, J. P. & Patraquim, P. Classification and function of small open reading frames. Nat Rev Mol Cell Biol 18, 575–589, doi:10.1038/nrm.2017.58 (2017).

75. Patraquim, P., Mumtaz, M. A. S., Pueyo, J. I., Aspden, J. L. & Couso, J.-P. Developmental regulation of canonical and small ORF translation from mRNAs. Genome biology 21, 128–128, doi:10.1186/s13059-020-02011-5 (2020).

76. Mirman, Z. & de Lange, T. 53BP1: a DSB escort. Genes & Development 34, 7–23, doi:10.1101/gad.333237.119 (2020).

77. Davuluri, R. V., Suzuki, Y., Sugano, S., Plass, C. & Huang, T. H.-M. The functional consequences of alternative promoter use in mammalian genomes. Trends in Genetics 24, 167–177 (2008).

78. Pakos-Zebrucka, K. et al. The integrated stress response. EMBO Rep 17, 1374–1395, doi:10.15252/embr.201642195 (2016).

79. Starck, S. R. et al. Translation from the 5′ untranslated region shapes the integrated stress response. Science 351, doi:10.1126/science.aad3867 (2016).

80. Steinberg, R. & Koch, H. G. The largely unexplored biology of small proteins in pro- and eukaryotes. The FEBS journal, doi:10.1111/febs.15845 (2021).

81. Zhang, P. et al. Relatively frequent switching of transcription start sites during cerebellar development. BMC Genomics 18, 461, doi:10.1186/s12864-017-3834-z (2017).

82. Zhang, H. et al. Genome-wide maps of ribosomal occupancy provide insights into adaptive evolution and regulatory roles of uORFs during Drosophila development. PLoS Biol 16, e2003903, doi:10.1371/journal.pbio.2003903 (2018).

83. Panier, S. & Boulton, S. J. Double-strand break repair: 53BP1 comes into focus. Nat Rev Mol Cell Biol 15, 7–18, doi:10.1038/nrm3719 (2014).

84. Kozak, M. Constraints on reinitiation of translation in mammals. Nucleic Acids Res 29, 5226–5232, doi:10.1093/nar/29.24.5226 (2001).

85. Luukkonen, B. G., Tan, W. & Schwartz, S. Efficiency of reinitiation of translation on human immunodeficiency virus type 1 mRNAs is determined by the length of the upstream open reading frame and by intercistronic distance. J.Virol. 69, 4086–4094 (1995).

86. McGeachy, A. M. & Ingolia, N. T. Starting too soon: upstream reading frames repress downstream translation. EMBO J 35, 699–700, doi:10.15252/embj.201693946 (2016).

87. Wethmar, K., Barbosa-Silva, A., Andrade-Navarro, M. A. & Leutz, A. uORFdb--a comprehensive literature database on eukaryotic uORF biology. Nucleic Acids Res 42, D60–67, doi:10.1093/nar/gkt952 (2014).

88. Pueyo, J. I. et al. Hemotin, a Regulator of Phagocytosis Encoded by a Small ORF and Conserved across Metazoans. PLOS Biology 14, e1002395, doi:10.1371/journal.pbio.1002395 (2016).

89. Bazzini, A. A. et al. Identification of small ORFs in vertebrates using ribosome footprinting and evolutionary conservation. 33, 981–993, doi:https://doi.org/10.1002/embj.201488411 (2014).

90. Kisselev, A. F. & Goldberg, A. L. Proteasome inhibitors: from research tools to drug candidates. Chemistry & Biology 8, 739–758, doi:https://doi.org/10.1016/S1074-5521(01)00056-4 (2001).

91. Dick, T. P. et al. Contribution of proteasomal beta-subunits to the cleavage of peptide substrates analyzed with yeast mutants. J Biol Chem 273, 25637–25646, doi:10.1074/jbc.273.40.25637 (1998).

92. Glickman, M. H. & Ciechanover, A. The ubiquitin-proteasome proteolytic pathway: destruction for the sake of construction. Physiological reviews 82, 373–428, doi:10.1152/physrev.00027.2001 (2002).

93. Schmidt, M., Hanna, J., Elsasser, S. & Finley, D. Proteasome-associated proteins: regulation of a proteolytic machine. Biological chemistry 386, 725–737, doi:10.1515/bc.2005.085 (2005).

94. Aiken, C. T., Kaake, R. M., Wang, X. & Huang, L. Oxidative stress-mediated regulation of proteasome complexes. Molecular & cellular proteomics : MCP 10, R110.006924–R006110.006924, doi:10.1074/mcp.M110.006924 (2011).

95. Dahlmann, B., Ruppert, T., Kuehn, L., Merforth, S. & Kloetzel, P. M. Different proteasome subtypes in a single tissue exhibit different enzymatic properties. J Mol Biol 303, 643–653, doi:10.1006/jmbi.2000.4185 (2000).

96. Pelletier, S. et al. Quantifying cross-tissue diversity in proteasome complexes by mass spectrometry. Mol Biosyst 6, 1450–1453, doi:10.1039/c004989a (2010).

97. Rivett, A. J. Intracellular distribution of proteasomes. Curr Opin Immunol 10, 110–114, doi:10.1016/s0952-7915(98)80040-x (1998).

98. Nederlof, P. M., Wang, H. R. & Baumeister, W. Nuclear localization signals of human and Thermoplasma proteasomal alpha subunits are functional in vitro. Proceedings of the National Academy of Sciences 92, 12060, doi:10.1073/pnas.92.26.12060 (1995).

99. Ogiso, Y., Tomida, A. & Tsuruo, T. Nuclear localization of proteasomes participates in stress-inducible resistance of solid tumor cells to topoisomerase II-directed drugs. Cancer Res 62, 5008–5012 (2002).

100. Kristiansen, M. et al. Disease-associated prion protein oligomers inhibit the 26S proteasome. Mol Cell 26, 175–188, doi:10.1016/j.molcel.2007.04.001 (2007).

101. Rubinsztein, D. C. The roles of intracellular protein-degradation pathways in neurodegeneration. Nature 443, 780–786, doi:10.1038/nature05291 (2006).

102. Quelle, D. E., Zindy, F., Ashmun, R. A. & Sherr, C. J. Alternative reading frames of the INK4a tumor suppressor gene encode two unrelated proteins capable of inducing cell cycle arrest. Cell 83, 993–1000, doi:10.1016/0092-8674(95)90214-7 (1995).

103. Klemke, M., Kehlenbach, R. H. & Huttner, W. B. Two overlapping reading frames in a single exon encode interacting proteins--a novel way of gene usage. Embo j 20, 3849–3860, doi:10.1093/emboj/20.14.3849 (2001).

104. Abramowitz, J., Grenet, D., Birnbaumer, M., Torres, H. N. & Birnbaumer, L. XLalphas, the extra-long form of the alpha-subunit of the Gs G protein, is significantly longer than suspected, and so is its companion Alex. Proc Natl Acad Sci U S A 101, 8366–8371, doi:10.1073/pnas.0308758101 (2004).

105. Lee, C. F., Lai, H. L., Lee, Y. C., Chien, C. L. & Chern, Y. The A2A adenosine receptor is a dual coding gene: a novel mechanism of gene usage and signal transduction. J Biol Chem 289, 1257–1270, doi:10.1074/jbc.M113.509059 (2014).

106. Liu, X. et al. Trip12 is an E3 ubiquitin ligase for USP7/HAUSP involved in the DNA damage response. 590, 4213–4222, doi:https://doi.org/10.1002/1873-3468.12471 (2016).

107. Legrand, N., Jaquier-Gubler, P. & Curran, J. The impact of the phosphomimetic eIF2αS/D on global translation, reinitiation and the integrated stress response is attenuated in N2a cells. Nucleic Acids Res 43, 8392–8404, doi:10.1093/nar/gkv827 (2015).

108. Terenin, I. M., Andreev, D. E., Dmitriev, S. E. & Shatsky, I. N. A novel mechanism of eukaryotic translation initiation that is neither m7G-cap-, nor IRES-dependent. Nucleic Acids Res 41, 1807–1816, doi:10.1093/nar/gks1282 (2013).

